# uSort-M: Scalable isolation of user-defined sequences from diverse pooled libraries

**DOI:** 10.64898/2026.01.12.699065

**Authors:** Micah B. Olivas, Patrick J. Almhjell, Lillian K. Brixi, Jack D. Shanahan, Polly M. Fordyce

## Abstract

Advances in high-throughput sequencing and computational protein design are expanding the catalog of known protein sequences far more rapidly than they can be functionally characterized. Functional characterization through biochemical and biophysical assays often requires isolating variants to profile them individually, a process which is often laborious and expensive. To address this, we developed <u>u</u>ser-defined <u>Sort</u>ed <u>M</u>utants (uSort-M), which can rapidly isolate and identify individual variants from widely available pools of genes by leveraging automated cell sorting and long-read sequencing technologies. To develop the uSort-M pipeline, we first parsed a 328-member scanning mutagenesis library of a 300-bp gene. Direct comparison between short-read and long-read sequencing demonstrated that both methods resolve variants with high fidelity, enabling recovery of 96% of desired library members by sorting eight 384-well plates at fivefold lower cost than traditional synthesis. After optimizing for long-read sequencing, we demonstrated uSort-M’s generalizability to complex libraries by parsing a sequence- and length-diverse 500-member library, recovering 88% of variants from shallow oversampling (<3-fold). Library recoveries could be accurately predicted by simulating library uniformity, transformation number, sorting efficiency, and per-base error rates, and we extrapolated these simulations to predict the sampling depth needed to process large libraries containing thousands of members. To facilitate adoption, these simulations are packaged alongside data analysis and workflow management tools in an open-source Python toolkit with an interactive dashboard. By implementing standard instrumentation in a generalizable workflow, uSort-M provides an efficient and cost-effective solution for large library generation, thereby removing a key barrier to large-scale protein functional characterization.

## Introduction

Advances in DNA sequencing and computational protein design are identifying new sequences of interest at unprecedented pace—from clinical variants and environmental orthologs to *de novo* designs and high-throughput screening hits.^1–7^ Experimentally characterizing these sequences requires the ability to rapidly and reliably generate hundreds to thousands of sequence-verified gene isolates.^3,8–11^ Despite advances in DNA synthesis, preparation of “parsed” libraries (where each gene is physically isolated and its sequence known) remains expensive, with per-sequence synthesis typically costing $10–100.

An alternative method is to prepare pooled DNA libraries and isolate single variants from the pool by stochastic sampling. This is a common approach in protein engineering, where degenerate codons are used to create a library of specified diversity from which clones are then randomly tested for activity, often without concomitant sequence-verification.^4,12–15^ In many cases, sequencing data is collected and mapped to functional measurements only for “hits”, yielding datasets without valuable negative data for building functional models. In engineering workflows, the number of clones tested is typically calculated based on the library size, with threefold oversampling predicted to return 95% of unique members in an unbiased library.^12,16,17^ This same sampling-based workflow can be applied to isolate and identify DNA variants by incorporating a sequencing step after parsing. However, existing methods typically require specific library designs, specialized equipment, and prescriptive sequencing pipelines.^8,9,18^ A general, accessible workflow for parsing pooled libraries into a physically isolated, sequence-verified format would remove a major experimental bottleneck and accelerate protein functional characterization at scale.

To address this need, we developed <u>u</u>ser-defined <u>Sort</u>ed <u>M</u>utants (uSort-M), a workflow that combines pooled DNA synthesis, widely accessible cell-sorting instruments, and recent advances in accurate, multiplexed long-read sequencing to generate isolated, verified DNA constructs assembled within desired plasmids in as few as seven days and for as low as $2.58 per variant (see **Methods** and **Supplemental Methods**). uSort-M eliminates time-consuming colony picking, whether by hand or via single-purpose robotics, and is fully agnostic to input library composition. Because uSort-M uses pooled inputs, the method benefits from continuously decreasing oligonucleotide synthesis costs and advances in pooled assembly methods.^19,20^ We describe the workflow, quantify how each step impacts library recovery efficiency, directly compare results from Nanopore sequencing to Illumina sequencing-by-synthesis for pooled amplicons, and use it to prepare two libraries of varying sequence and length diversity. We further provide tools for estimating recovery, simulating effects of critical variables like library bias, and demultiplexing sequencing reads to identify expected library members. We anticipate that this workflow and accompanying tools will be broadly useful across library types and substantially reduce the time and cost of creating defined collections of isolated constructs.

## Results

### Overview of the uSort-M pipeline

The uSort-M pipeline rapidly isolates and sequences individual DNA variants from a pre-synthesized oligo pool via stochastic sampling, yielding a final parsed and sequence-validated library (**Figure 1A**). uSort-M involves: 1) assembling a pooled library into a plasmid vector of choice; 2) transforming *Escherichia coli* (*E. coli*) cells with the pooled library and culturing overnight; 3) sorting individual transformed *E. coli* into wells of 384-well plates via FACS and culturing overnight under antibiotic selection; 4) amplifying DNA templates and appending plate/well-based barcodes via colony PCR; and 5) pooling PCR amplicons prior to multiplexed sequencing (**Figure 1B**). Each sequencing read contains the sequence of the variant and a barcode pair specifying the plate/well-based position; the many reads that are mapped to each well are then used to identify the DNA variant within that well. Desired library members can be selected from these plates and organized for downstream applications.

**Figure 1.**
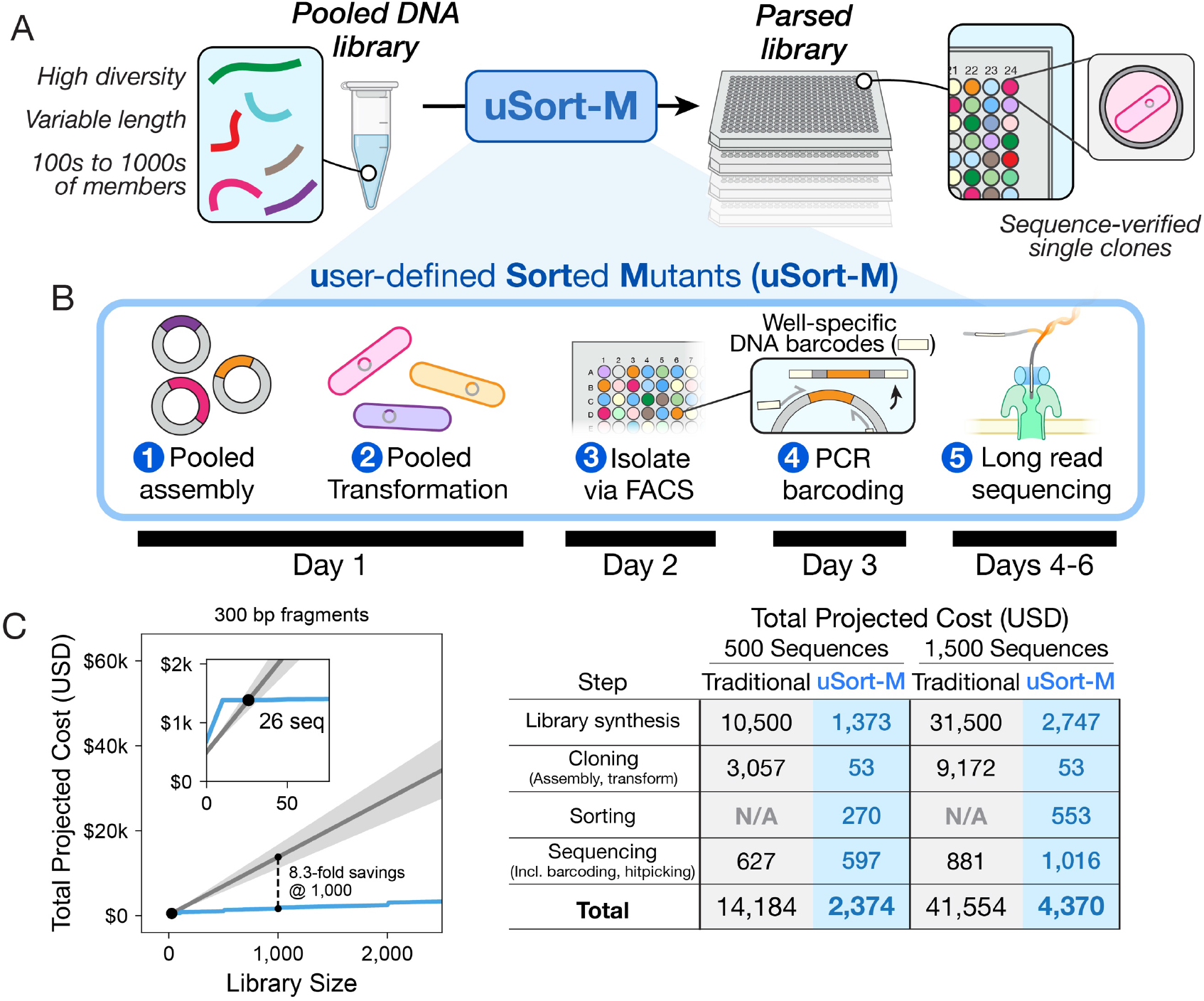
Overview of the uSort-M pipeline. **A.** uSort-M converts an arbitrary pooled library of DNA variants into a “parsed” library where each variant is isolated in its own well with validated sequence information. **B**. uSort-M isolates single plasmid variants within a bacterial host via: (1) pooled assembly, (2) transformation, (3) sorting individual bacteria into wells via high-throughput cell sorting (FACS), (4) amplifying and barcoding DNA amplicons from cultured clones via PCR, and then (5) pooling amplicons for multiplexed long-read sequencing that associates plate/well-specific barcodes with DNA amplicons. **C**. Projected costs of traditional library preparation *vs*. uSort-M. *Left*: cost as a function of library size for traditional synthesis methods (light grey shading indicates estimated minimum and maximum costs; grey line indicates mean) *vs*. uSort-M (blue line) for libraries of 300 bp fragments. Dashed black line indicates 8.3-fold savings for uSort-M vs traditional synthesis. *Right*: table indicating the estimated cost of individual steps required for traditional gene synthesis or uSort-M for 500- and 1,500-variant libraries of 300 bp fragments.

uSort-M provides two main advantages relative to previously published techniques. First, most prior approaches isolate single transformed clones from a pool by either manual colony picking (which is labor-intensive) or using a specialized and expensive colony-picking robot.^8,12^ uSort-M employs a fluorescence-activated cell sorting (FACS) instrument to isolate single *E. coli* cells directly from liquid culture. Commercial and widely available FACS instruments can sort single bacteria at high rates (∼70 wells per minute in single-cell sorting mode) even in the absence of a fluorescent signal, making this a highly general approach.^21–24^ Second, most prior approaches have sequenced amplicons using Illumina-based sequencing that limits read lengths to ∼300–700 nucleotides (paired-end 150- or 600-cycle MiSeq v3 kits).^8,14^ The need to associate DNA barcodes with sequence information throughout an amplicon either limits the maximum size of the region of interest to ∼300 bases or requires additional transposase-based fragmentation. uSort-M instead implements Nanopore sequencing, which returns reads that can span >25 kilobases,^25,26^ has been validated for barcode-based demultiplexing of pooled libraries,^15,27^ and is available through several commercial providers with turnaround times ranging from overnight to a few days for tens of thousands or multi-gigabase (Gb) read depths, respectively.

uSort-M becomes more cost-effective than traditional gene fragment synthesis for 300-bp libraries with as few as 26 constructs (**Figure 1C** and **Supporting Information, Figure S1** and **Table S1**). As library size increases, savings grow substantially, reaching ∼80% cost reduction at 1,000 variants.

### Equivalent performance of short- and long-read sequencing on an initial uSort-M library

To evaluate uSort-M performance on a realistic library generation task, we sought to directly compare results from Illumina’s short-read sequencing and Nanopore’s long-read sequencing. Illumina has been the standard for deep amplicon sequencing applications because of its high fidelity and throughput^28^, providing efficient and robust demultiplexing of pooled libraries. We ordered a 328-variant pooled oligonucleotide library encoding four different codon substitutions at each of 85 residue positions (excluding wild-type amino acids) within the small human enzyme acylphosphatase-2 (hAcyP2)^29^ (**Figure 2A**). hAcyP2’s small size (∼100 amino acids) allowed us to encode the entire structured region within a ∼250-bp oligonucleotide, simplifying synthesis (**Supporting Information, Table S2**). After appending ∼350 bp of DNA sequence required for plate/well position barcoding and Illumina sequencing (see below), each amplicon remained ≤650 bp and could be sequenced using both a MiSeq and Nanopore to enable direct comparison on the same input library (**Supporting Information, Figure S2**).

**Figure 2.**
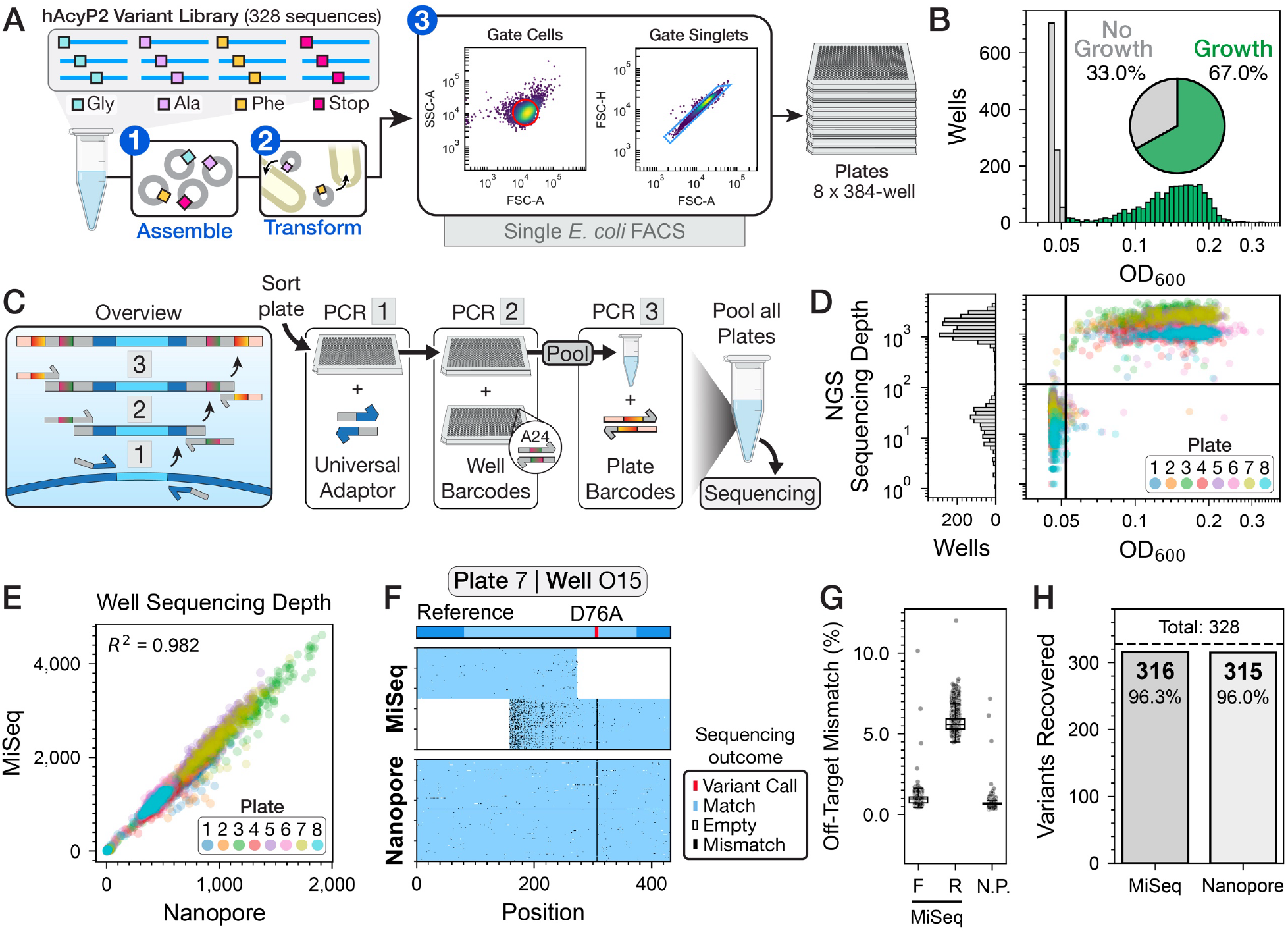
Efficient recovery of a 328-variant scanning library with uSort-M. **A.** Schematic illustrating 328-variant human acylphosphatase 2 (hAcyP) library processing, including: (1) assembly, (2) transformation, and (3) FACS sorting. Left scatter plot: selection of cell events by gating the distribution of side scatter area (SSC-A) *vs*. forward scatter area (FSC-A) values. Right scatter plot: selection of singlets from gating the distribution of forward scatter height (FSC-H) *vs*. FSC-A. **B**. Histogram of the number of wells as a function of measured OD_600_ values. Wells with bacterial growth (green bars representing 2,057 wells) were defined as having OD_600_ > 0.052 (grey vertical line). Inset pie chart indicates fraction of wells with (green, 67%) and without (grey, 33%) growth. **C**. Plate-well barcoding scheme for multiplexed sequencing of variant DNA in each well. A sequence of PCR steps encodes the location of the source well (a two-step, one-pot PCR; steps 1 & 2) and plate (step 3) for each read. **D**. Number of demultiplexed MiSeq reads (sequencing depth) *vs*. well OD_600_; marker color indicates plate of origin; vertical and horizontal lines are shown for OD_600_ = 0.052 and sequencing depth = 100, respectively. Histogram (left) indicates number of wells with a given read depth. **E**. Scatter plot comparing number of reads per well from demultiplexed MiSeq data *vs*. Nanopore data; annotation specifies coefficient of determination (*r*^*2*^). **F**. Read alignment plots from a representative well comparing paired-end MiSeq data and long-read Nanopore data. Sequence positions are aligned along the x-axes of each plot, with bases matching the reference colored in light blue, mismatches in black, and empty positions in white. *Top*: read schematic showing aligned positions corresponding to the hAcyP2 WT reference sequence (light blue), called variant for the given well (red), and DNA appended during indexing (dark blue). *Middle*: Subsampled paired-end MiSeq reads ordered by orientation, with forward reads on top and reverse reads on bottom. *Bottom*: Subsampled Nanopore reads. **G**. Quantified fidelity across all reads from all wells, defined as the frequency of finding a mismatch outside of the called variant bases. Each point shows the fidelity of a single read, boxes denote the median, 25^th^, and 75^th^ percentiles of the distribution, and error bars represent the standard deviation. (N.P.: Oxford Nanopore.) **H**. Number of variants recovered for the 328-member library from MiSeq or Nanopore data.

We amplified the oligonucleotide pool using a universal flanking primer set and then assembled it into a destination vector containing an ampicillin resistance marker via pooled Golden Gate cloning (**Figure 2A**, steps 1 and 2).^30^ After transformation, plating, and overnight growth, we obtained 5,250 transformants, representing a 16-fold oversampling of the 328-member library. By Poisson sampling statistics from a 328-member population with replacement, this level of oversampling represents an expected >99.9% completeness of the library in the sampled transformants^12,16,17^ assuming uniform representation of each DNA variant (see **Methods** for calculations).

After culturing the pool of transformed cells overnight, we used a BD FACSAria II to sort individual bacteria into 384-well plates (**Figure 2A**, step 3). Gating based on forward and side-scattering signals discriminated bacteria from other debris in the absence of a fluorescent signal; an additional gate based on the height and area of the forward scattering signals identified single bacterial cells. Anticipating some loss at each subsequent stage of the pipeline, we sorted eight 384-well plates over ∼2 hours of instrument time (3,072 wells total; 9.4-fold oversampling and >99.9% expected completeness, **Figure 2A**). After sorting and culturing overnight, 2,057 of 3,072 wells showed growth (as assessed by quantifying optical density at 600 nm), equivalent to 67% efficiency, 6.3-fold oversampling, and 99.8% expected completeness (**Figure 2B** and **Supporting Information, Figure S3** and **Table S3)**. Because plates were cultured without shaking, colonies were visible at the bottom of many wells with growth (others appeared suspended), which made it possible to manually examine or image plates to detect doublets ahead of sequencing (**Supporting Information, Figure S4**).

To enable pooled sequencing of amplicons while preserving information about their well location, we used colony PCR to amplify DNA templates directly from the cultures and append primer barcodes encoding plate and well positions. Illumina’s Nextera indices served as plate-specific outer barcodes (**Supporting Information, Table S4**) and evSeq indices served as well-specific inner barcodes according to a standardized mapping scheme^14^ (**Figures 2C and S2**, see **Methods** and **Supporting Information**). The Nextera barcoding primers also appended the i5 and i7 adapter sequences required to perform Illumina sequencing on a MiSeq. Oxford Nanopore sequencing directly ligates its required adapters to DNA molecules prior to sequencing, allowing us to sequence the same exact pool of DNA via both methods and directly compare outcomes. The full cost of barcoding variants was ∼$0.25 per well, or ∼$100 per 384-well plate (**Supporting Information, Table S1**).

After obtaining sequencing data from both methods, we examined how individual reads were mapped to plate-well positions using their respective demultiplexing pipelines. As expected, Illumina MiSeq reads were efficiently mapped to individual wells using existing methods designed for this purpose (**Methods**, see **Supporting Information, Table S5** for compiled demultiplexing results). We obtained 5.54 M paired-end reads containing the correct Nextera barcodes using Illumina’s bcl2fastq demultiplexing software, which assigned reads to their respective plates; we then used the evSeq demultiplexing software^14^ to further assign the inner barcodes to their respective wells within each plate. Demultiplexing via evSeq assigned 3.45 M reads to wells, with the remaining reads failing due to low read quality or the absence of an expected barcode pair. There were two clear distributions for wells with high (>100 reads) vs. low sequencing depth, which corresponded to wells with high vs. low bacterial growth in the input culture (**Figure 2D**). Overall, 98% (2,017/2,057) of wells with high bacterial growth after sorting were assigned >100 reads, confirming robust amplification of DNA templates directly from bacterial culture. A lack of reads from wells without growth additionally confirmed minimal contamination or index hopping during multiplexed sequencing library preparation.

Equivalent demultiplexing results were obtained using the Nanopore pipeline. Multiplexed Nanopore sequencing typically uses 24-bp barcodes to compensate for a higher expected level of base-calling noise^15,26,28,31,32^, which is longer than either the 12-bp outer Nextera barcode or 7-bp inner evSeq barcode used here. Nevertheless, Nanopore sequencing reads containing these shorter barcodes were readily demultiplexed by Oxford Nanopore’s Dorado software^33^ (**Methods**), assigning 2.28 M of 4.31 M reads to plates and 1.47 M of those 2.28 M reads to specific wells. Per-well sequencing depths were ∼2-fold lower than, but highly correlated with, MiSeq-based results across all 8 plates (**Figure 2E**, *r*^2^ = 0.98, slope = 0.42). Despite higher reported sequencing error rates for Nanopore sequencing and relatively short 7-bp barcodes, the MiSeq+evSeq and Nanopore+Dorado pipelines had comparable rates of assigning plate reads to wells (62.2% vs. 64.4%, **Supporting Information, Table S5**) and Nanopore sequencing returned >100 reads for 2,005 of the 2,057 wells with growth (97%). Nanopore reads were no more likely than MiSeq reads to be assigned to wells lacking obvious growth (**Supporting Information, Figure S5**), suggesting that random miscalling or misassignment of the shorter barcodes did not occur.

We next aligned well-specific reads to the reference hAcyP2 sequence to determine the variant contained within each well. For the MiSeq data, we continued to use the established evSeq software to determine the identity of each well; for the Nanopore data, we used minimap2^34^ to align reads to the reference sequence and custom software to determine mismatched bases and confirm final variant calls using the samtools consensus software^35^ (**Methods**). Across the 2,005 wells that had >100 Nanopore and MiSeq reads, 1,676 (83.6%) had identical variant calls across both platforms. Of the 329 that did not match, 284 (86%) of these were multiclonal. Of the remaining 45 wells, all but one contained indels. These indels were directly detected from Nanopore sequencing by samtools consensus but were filtered out during the evSeq variant calling pipeline to a low-but-nonzero number (<100) of spurious indel-filtered reads. The single remaining well that passed all quality-control thresholds but did not match was caused by an instance where the Nanopore pipeline identified mismatches in the same codons as the MiSeq pipeline but was unable to resolve their specific identities. Overall, Nanopore errors were randomly distributed across reads such that true variation was easy to identify (**Figure 2F**). By contrast, Illumina read errors were similarly distributed in the forward reads but highly concentrated near the end of the reverse reads (**Figure 2F,G**). Both sequencing methods recovered variants at similar frequencies across positions in the hAcyP2 amplicon, with no evidence of positional bias affecting variant recovery (**Supporting Information, Figure S6**).

Finally, we assessed the number of desired variants recovered by MiSeq and Nanopore sequencing, respectively. For the 2,017 wells with >100 MiSeq reads, 1,375 (68.2%) contained desired single-mutant sequences as their most abundant variant. This suggests that ∼32% of wells contained off-target variation (e.g., multiple mutations or indels). For our 300-mer pooled oligos with a synthesis error rate of 1:3000 bases, we expected 10% of oligos to contain errors, suggesting that PCR amplification of the ssDNA pool, Golden Gate assembly, and *E. coli* processes during transformation contributed the remaining ∼22% of off-target variation. Of these 1,375 samples, 79 had low clonal purity (<90% of reads indicated the same variant; **Supporting Information, Figure S7**), leaving a total of 1,296 in-library monoclonal wells (64.2%, 4-fold oversampling of the 328-member library). Nanopore variant calling results were similar, with 1,293 (64.5%) of 2,005 wells with >100 reads containing high-confidence monoclonal single-mutant sequences. From these sampling numbers, we expected ∼98% coverage of the 328-member library. Consistent with this prediction, we identified 316 (96.3% completeness) and 315 (96.0% completeness) unique variants using the MiSeq and Nanopore pipelines, respectively (**Figure 2H**). Overall, Nanopore offered lower cost and higher flexibility while returning similar information, even when using libraries with sequence features optimized for Illumina sequencing.

### Simulations recapitulate experimental results and reveal factors that influence library recovery

Standard sampling statistics predict library recovery without considering the potential impact of inefficiencies throughout the uSort-M pipeline, potentially explaining why our measured recovery rate (96%) was slightly below the predicted rate (98%). To estimate the number of wells that must be sorted and sequenced for a given library coverage under real-world conditions, we simulated how inefficiencies in DNA synthesis and assembly, bacterial transformation, sorting and outgrowth, and sequencing library preparation impact variant recovery. Each simulation considers five primary user-defined quantities: (1) **library skew**, or the fold-difference in relative abundance between the most- (90^th^ percentile) and least- (10^th^ percentile) represented sequences^36^; (2) **off-target variation**, or the fraction of amplicons that do not match the target library due to errors during DNA synthesis, plasmid assembly, transformation, and outgrowth, (3) **transformation scale**, or the ratio of the number of transformed cells to the number of unique library variants, (4) **sorting and outgrowth efficiency**, or the fraction of wells that show growth, and (5) **PCR failure rate**, or the fraction of wells with growth that do not yield product after colony PCR (**Figure 3A**).

**Figure 3.**
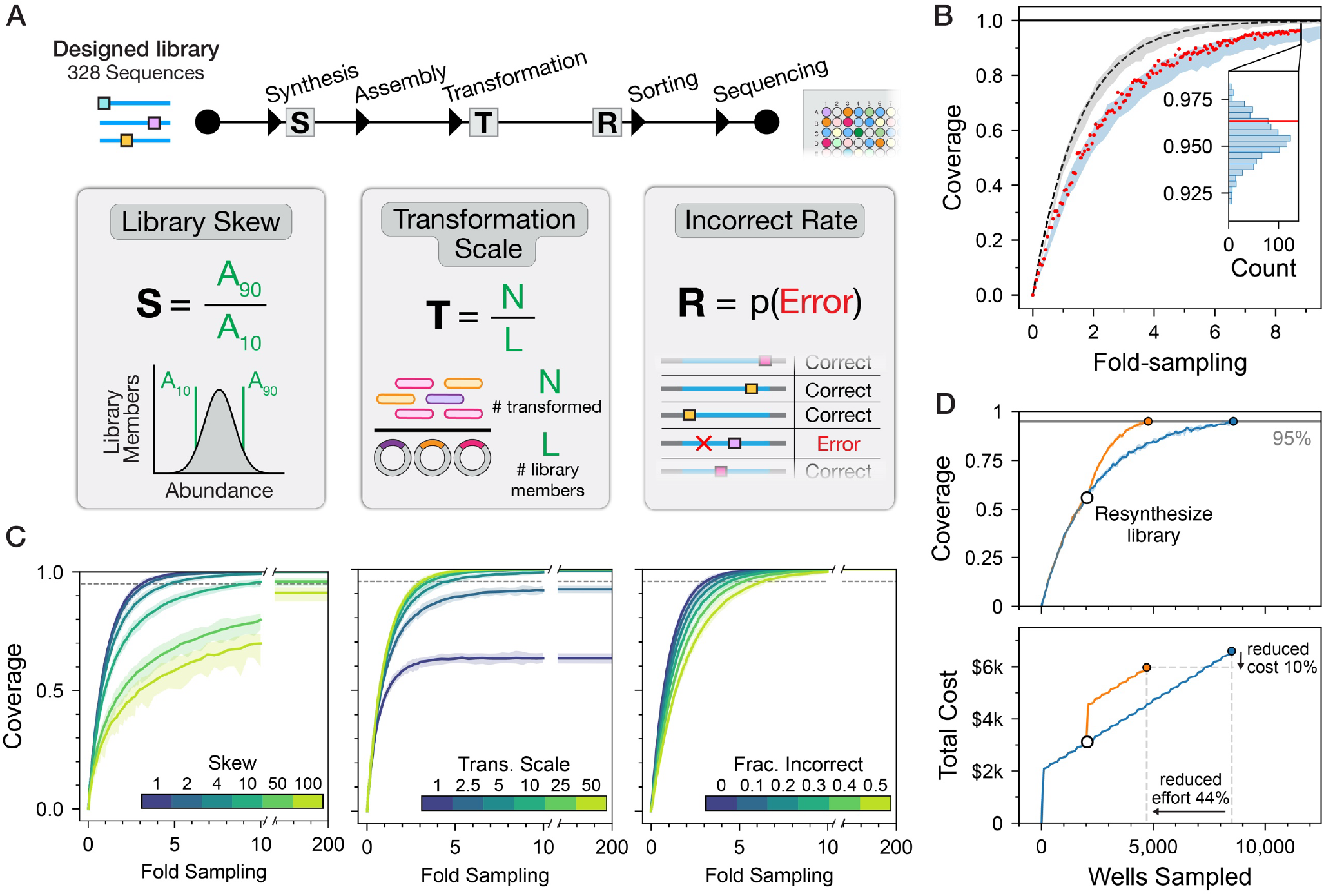
Simulations demonstrating the effects of uSort-M parameters on library coverage. **A.** Simulation pipeline illustrating key parameters from each step in the uSort-M workflow and how they are defined. **B**. Simulated coverage as a function of fold-sampling for a 328-member library, where the number of wells sorted is equal to the product of the library size and fold-sampling. Red points represent bootstrap downsampling from the collected hAcyP2 library sequencing data for each fold-sampling value; blue shaded area shows 99% confidence interval from 100 independent simulations using known parameters of the experiment (skew = 2, off-target variation = 0.35, transformation scale = 5,250/328, sorting efficiency = 0.67, PCR failure rate = 0.025). Grey shaded area shows 99% confidence interval from 100 independent numerical simulations assuming a ‘perfect’ input library (skew = 1, off-target variation = 0, transformation scale = 100, same sorting and PCR parameters); dashed black line indicates the analytical solution to sampling from this population. Inset shows simulated coverage distribution at 3,072 sorted wells in blue; red line indicates the observed coverage (0.963, 316 variants). **C**. Coverage *vs*. fold-sampling plots for a 1000-member library with varying skew, off-target variation, and transformation scale, holding non-varied parameters at ‘realistic’ values of 1, 0, and 50, respectively. The x-axis is discontinuous between 10x sampling and >380x sampling to illustrate where the curves plateau. **D**. Coverage (top) and cost (bottom) *vs*. wells sampled for a 1,000-member library after either exhaustive sampling (blue curve) or sampling with a ‘targeted resynthesis’ step (orange curve). In ‘targeted resynthesis’, a set number of wells are sorted and sequenced to a lower coverage value, and then the remaining unsampled members of the library are reordered as a new, smaller library.

To test if these parameters are sufficient to accurately explain our observed uSort-M recovery rate, we simulated recovery based on the known parameter values from the initial hAcyP2 scanning library (a library size of 328, a library skew of 2, a total of 5,250 transformants, 35% off-target variation, 67% sorting outgrowth efficiency, and a 2.5% PCR failure rate; **Supporting Information, Figure S8**). These simulations predicted a median recovery of 96% at 3,072 wells, in agreement with the observed median recovery of 96% (**Figure 3B, inset**). Predicted recoveries across all fold-sampling values were also in excellent agreement with down-sampling simulations that mimicked picking fewer wells from the hAcyP2 experiment, especially when compared to unbiased sampling conditions (**Figure 3B, full curves**), further establishing that the uSort-M process was well-approximated by these simulations.

Next, we systematically varied library skew, transformation scale, and off-target variation for a simulated 1000-member library to assess how each parameter affects coverage (**Figure 3C** and **Supporting Information, Table S9**). The simulations revealed that differences in library skew have the largest impact on recovery: at 1-, 10-, and 50-fold skew, achieving 90% coverage requires oversampling by 2.4, 5.8, and 27.6-fold, respectively. For high (50-fold) skew, even 200-fold oversampling (200,000 wells, in this case) recovers only 95% of variants, as rare variants must compete against substantially more abundant sequences during sampling. As shown in **Figure 3C**, transformation scale sets an upper bound on the coverage that can be achieved, as variants that don’t appear in the initial transformed pool can never be recovered. At a transformation scale of 1 (the number of transformants is equal to the library size), only 63% of variants from a perfectly uniform library are expected to be present in the transformed cells (again calculated using Poisson sampling statistics). Increasing transformation scale to 2.5 increases this to ∼90% recovery (regardless of wells sampled) and transformation scales of 5 or higher ensure near-complete representation for uniform libraries. Off-target variation acts as a constant tax on sampling efficiency by introducing “wasted” samples in which wells contain incorrect sequences. Off-target variation typically stems from errors that occur during DNA synthesis (Twist Biosciences reports a 1:3000 bp error rate) or pooled library preparation (e.g., template switching) and can range from 10– 40%.^8^ At a 30% off-target rate (near the ∼35% observed in the hAcyP2 library), achieving 90% coverage requires 4-fold oversampling (compared to 2.5-fold with perfect fidelity). Although **Figure 3C** varies each parameter independently, in practice their effects compound, making prospective simulation particularly important for experimental design. The full data and model can be found at https://github.com/FordyceLab/usortm.

### Charting the efficient recovery of large and highly skewed libraries with iterative resampling

For large or highly skewed libraries, sampling redundancy increases substantially as previously isolated variants are re-sampled (**Figure 3C**). As a result, attaining high library coverage via a single round of sampling can require sorting tens of thousands of wells across many dozens of 384-well plates. In these cases, iterative resampling with targeted resynthesis can dramatically increase efficiency. In this scheme, an initial round of modest oversampling (e.g., 2-fold) is applied to efficiently sample unique variants in a regime where the rate of unique variant recovery per sampled well is nearly linear. After this initial stage, missing variants are then resynthesized as a new oligo pool and used for a second round of sampling. For a 1,000-member library with 2-fold skew and 30% off-target variation (**Figure 3D**), a single-pass approach would require sorting 8,569 wells (∼8.5-fold oversampling) to achieve 95% coverage. By contrast, one round of resampling at 2-fold coverage would achieve the same coverage with 4,743 total sorted wells—a 44% reduction. Incorporating a resynthesis step into our Monte Carlo simulation, we found an overall cost of ∼$6,000 at 2-fold resampling (compared with $6,600 for a single-pass approach). By simulating resynthesis across various initial sampling depths, we identified approximately 2-fold oversampling as the optimal trigger point for resynthesis of large libraries (**Supporting Information, Figure S9**). This yields considerable savings for large libraries: for a 3,000-member library, resynthesizing the library at 2-fold oversampling yields more than 20% cost savings and ∼10,000 fewer total wells sorted compared to a single-pass workflow (**Supporting Information, Figure S9**).

### A uSort-M pipeline optimized for long-read sequencing enables robust recovery of a diverse, length-variable library

The hAcyP2 library leveraged the evSeq barcoding scheme to enable a direct comparison of Nanopore and MiSeq results (**Figure 2**). Based on the favorable results from Nanopore, which provides a more convenient and flexible sequencing strategy, we opted to optimize the uSort-M pipeline for this approach. We then evaluated its performance on the generation of a library with greater sequence and length diversity than our hAcyP2 scanning library. This new library was comprised of 500 intrinsically disordered regions (IDRs) from 223 distinct human protein domains that were each represented by 1–45 unique sequence variants ranging from 30–146 residues in length (90–438 bp coding sequence) (**Figure 4A**). Pairwise sequence distances spanned near-identical variants to highly divergent sequences, providing a distinct challenge from the low diversity, constant-length hAcyP2 scanning library.

**Figure 4.**
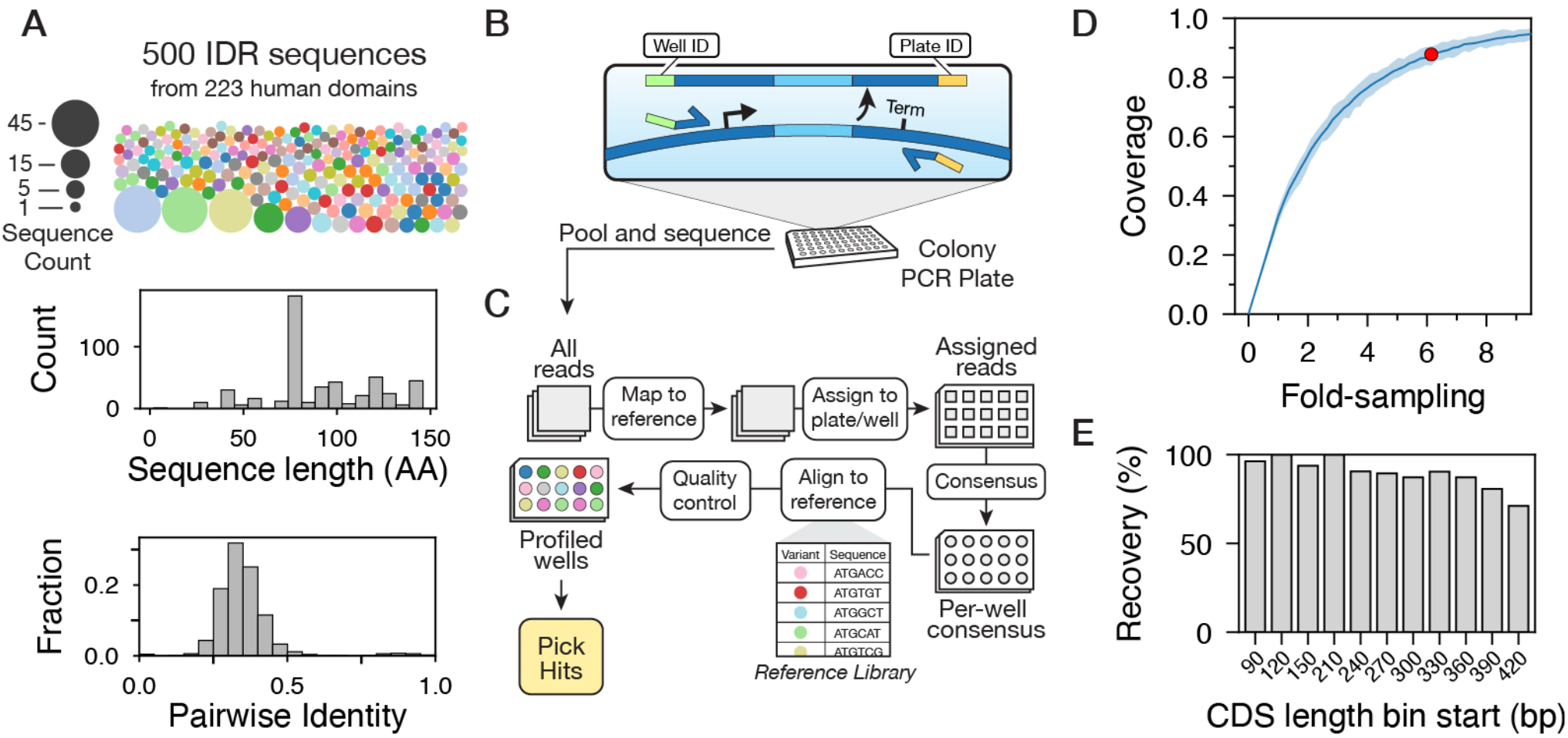
Recovery of a diverse, length-variable library. **A.** Composition of the 500-member IDR library. Top: Each “reference” domain is shown as a circle with its size corresponding to the number of different variants (e.g., substitution mutants) of that reference within the library. Bottom: Histograms of sequence length (sequence counts *vs*. number of amino acids) and pairwise identity (fraction of sequences *vs*. pairwise identity). **B**. Simplified barcoding scheme. The forward primer encodes a well-specific barcode (green) and the reverse primer encodes a plate-specific barcode (yellow). Primers bind in a common region in the vector (dark blue) outside of the variable region of interest (light blue). **C**. Processing workflow for nanopore sequencing data. **D**. Coverage plot, same as in Figure 3B. The red point shows obtained coverage and fold-sampling for this library. Blue shaded area shows 99% confidence interval from 100 independent simulations using known or expected parameters of the experiment (skew = 2, off-target variation = 0.3, transformation scale = 87, sorting efficiency = 0.67, PCR failure rate = 0.05). **E**. Bar plot of recovery rate across CDS length (bin width = 30 bp; the 180-210 bp bin is omitted due to a lack of sequences in that length range).

Each IDR was cloned as a SNAP-IDR-eGFP fusion within an expression vector containing a T7 promoter and terminator (**Supporting Information, Figure S10** and **Supplementary Data**). After pooled IDR library cloning, we transformed *E. coli* and sorted single bacteria into eight 384-well plates (3,072 wells total; targeting ∼6-fold oversampling). After overnight outgrowth in LB, 63.3% (1,946/3,072) of wells showed growth, representing 3.9-fold oversampling. We barcoded these IDR amplicons with well- and plate-based indices via the LevSeq barcoding scheme^15^, a long-read-adapted version of evSeq (**Figure 4B**). In LevSeq, the forward and reverse primers encode well and plate positions, respectively, and bind common flanking regions within the destination vector outside of the variable region (here, the T7 promoter and T7 terminator). This scheme thus generates up to 96^2^=9,216 unique plate/well barcode pairs from two 96-well plates of primers (one forward and one reverse) and barcodes amplicons from any sorted, transformed *E. coli* in a single PCR step.

In parallel, we expanded the demultiplexing pipeline for compatibility with sequence-diverse libraries (**Figure 4C**). The initial alignment step with minimap2^34^ aligned reads to the 5’ and 3’ sequences of the transformation vector immediately flanking the variable region, confining consensus generation to a window defined by the surrounding constant regions. By anchoring on invariant flanking sequences rather than the variable region itself, this approach accommodates inserts of arbitrary sequence and length, including libraries with no shared homology between members. We implemented this pipeline in a Nanopore-optimized software package that takes as inputs a FASTA file with all library sequences and the multiplexed FASTQ sequencing data and returns a user-friendly, interactive reporting summary that identifies wells containing desired library members and enables well-specific analysis (available at https://github.com/FordyceLab/usortm) (**Methods**).

Colony PCR with LevSeq primers and multiplexed Nanopore sequencing yielded 1.1M reads, of which 52.6% were successfully demultiplexed to individual wells (compared to 34.1% from the initial implementation with evSeq’s shorter, dual-index barcodes; **Supporting Information, Table S5**). Variant calling identified 1,348 wells containing high-confidence monoclonal sequences (≥90% of reads supporting a single variant call) corresponding to library members, equivalent to ∼2.7-fold sampling. From a single round of sorting and processing eight 384-well plates, we recovered 439 of 500 unique IDRs (87.8% completeness), agreeing with sampling simulations based on an expectation of 2-fold library skew (∼87%; **Figure 4D**). Recovery rate was lower for variants with longer coding sequences, likely due to synthesis errors accumulating at longer sequence lengths (see **Figure 4E**).

After a single round of resynthesis in which we ordered 39 missing genes as isolated eBlocks from Integrated DNA Technologies and used the uSort-M sequencing pipeline to barcode and analyze four clones per gene (156 wells total), we increased variant library coverage to 95.6%. These results establish that uSort-M, with single-step LevSeq barcoding and flank-anchored demultiplexing, can efficiently parse libraries with substantial diversity in sequence identity, amplicon length, and domain composition.

## Conclusions

uSort-M provides a scalable and cost-efficient path toward generating large libraries of isolated genes. Pairing widely accessible FACS instruments for clone isolation with commercial long-read nanopore sequencing bypasses the need for specialized robotics and circumvents amplicon length and composition limitations inherent to short-read sequencing-by-synthesis. uSort-M is fully agnostic to the composition of the input pool: any method that yields transformable plasmid DNA (e.g., assembly of tiled variable regions or entire genes^20,37,38^, pooled site-directed^8^ or site-saturation^12,13^ mutagenesis, error prone PCR^39^) can be parsed using this workflow, as demonstrated with our highly diverse library of IDRs.

Sampling simulations revealed that library skew exerts the strongest influence on recovery efficiency, with substantially greater oversampling required to achieve high coverage of rare variants. Commercial pooled oligonucleotide synthesis (e.g., Twist Biosciences, IDT) provides highly uniform libraries but is commonly limited to sequences shorter than 350 bp. Recent advances in pooled assembly are extending accessible fragment lengths by engineering orthogonal, pairwise interactions in diverse pools.^20,40^ As these methods make longer pooled libraries more broadly accessible, library uniformity will increasingly govern parsing efficiency—making prospective simulation of recovery, enabled by uSort-M, a practical necessity for experimental design.

For either large or highly skewed libraries, iterative resampling with targeted resynthesis reduces total sorting effort by ∼50% compared to single-pass, exhaustive oversampling approaches. This iterative strategy trades an increased cost of pooled library synthesis against reduced costs and effort of sorting and barcoding. While the total costs are typically only marginally lower, the total effort and plate storage requirements are greatly reduced. The simulation framework and demultiplexing tools provided at https://github.com/FordyceLab/usortm allow users to predict the required effort and expected coverage for a given library composition before beginning experiments, enabling optimization of the trade-off between sorting effort and resynthesis cost.

In future work, the time and costs required for library generation via uSort-M could be improved by increasing sorting efficiency. Currently, we reliably culture cells and recover amplicons from 60–70% of wells across all tested instruments (e.g., Sony SH800 and BD FACSAria II) and conditions (e.g., 70- or 100-µm nozzles, stationary phase or log phase growth, viability dye or scattering only). This number is ∼2-fold higher than expected for Poisson loading of highly dilute cultures (“limiting dilution”)^41^, but lower than what can be achieved by colony picking. Optimization of sorting and recovery could decrease the number of wells that must be processed, yielding substantial savings in time and reagent costs. As an additional improvement, constructs containing a fluorescent protein tag could be directly sorted based on fluorescence, eliminating a need for overnight culture on selective media to identify successful transformants. We anticipate that this workflow will be broadly useful for generating defined collections of protein variants for downstream biochemical characterization, accelerating efforts to link sequence to function at scale.

## Methods

### Cost analysis

Costs for traditional gene synthesis were calculated using pricing from three commercial gene fragment products: IDT gBlocks ($0.09/bp), IDT eBlocks ($0.07/bp), and Twist Gene Fragments ($0.07/bp). Costs for the full uSort-M workflow were calculated as the sum of pooled synthesis, cloning, sorting, barcoding, and sequencing. Costs are listed in **Supporting Information, Table S1**. Pooled library inputs were synthesized using Twist Oligo Pools (pricing available at https://www.twistbioscience.com/products/oligopools and in **Supporting Information, Table S2**). Cloning costs were calculated assuming a single-insert, 100-µL Golden Gate Assembly reaction into the user’s destination vector, followed by transformation into chemically competent cells. Assembly reactions were projected to consume 50 µL of 2X NEBridge Golden Gate Assembly Master Mix (NEB E1601L), corresponding to $53.60 in reagent costs. Transformation reagent costs constituted $3.13 for 25 µL of NEB 5-alpha Ultracompetent cells (NEB C2987H). Sorting costs were calculated based on instrument and operator time at a combined core facility rate of $135/hr ($70/hr for instrument time, $65/hr for operator time). At approximately 5 minutes per 384-well plate, sorting cost $11.25 per plate. Sorted cells were collected in 384-well plates (Greiner 781091). Barcoding costs, including primers and PCR reagents, totaled $97.73 per 384-well plate. Sequencing costs were calculated using Plasmidsaurus Custom Sequencing pricing, which starts at $500 for 1 gigabase of sequencing and increases by $50 for each added gigabase (as of 2025; https://www.plasmidsaurus.com/custom). Anticipated sequencing depth was fixed at 100-fold coverage of the amplicon library; this target depth was used to compute the required sequencing volume for each library size.

The minimum cost per variant ($2.58) represents an optimized scenario using short oligonucleotide substitutions (30 bp) assembled into a 2,000 bp wild-type ORF, assuming minimal library skew (≤2-fold), moderate sorting efficiency (∼67%), and moderate oversampling (6x). Actual costs vary with library size, fragment length, and complexity (see **Figure 1C** and **Supporting Information, Supplemental Methods** for detailed justification and cost projections).

### Scanning library design and synthesis

Scanning libraries were based on a sequence of human acylphosphatase 2 (hAcyP2; UniProt ID: P14621) that was codon-optimized for expression in *E. coli*. Each in-frame codon was mutated to code for Gly (`GGC`), Ala (`GCG`), Phe (`TTT`), or a stop codon (`TAG`), excluding synonymous mutations, from position 9 to 93. A single-stranded oligonucleotide pool containing each sequence flanked by BsaI restriction sequences and terminal primer adapter sequences was ordered from Twist Biosciences. The wild-type hAcyP2 DNA sequence and destination vector sequence are available in the **Supporting Information**.

### AcyP library cloning and transformation

The oligonucleotide pool was resuspended to 20 ng/µL in DI water. The ssDNA library was converted to dsDNA following the manufacturer’s protocol: 0.5 µL of the sample was used to perform PCR using a Roche KAPA HiFi HotStart ReadyMix PCR Kit (Roche KK2602; final DNA concentration 0.4 ng/µL) and subjected to the following thermal cycle: 95 °C, 3 min; 12 × [98 °C, 20 s; 62 °C, 15 s; 72 °C, 15 s]; 1 min, 72 °C.

Amplicon products were analyzed and isolated by gel electrophoresis and recovery (Zymo D4001). Purified DNA was assembled into a plasmid destination vector (see **Supporting Information**) via a Golden Gate^30^ reaction using NEBridge BsaI-HFv2 Golden Gate master mix (NEB E1601S). Assembly reactions were used directly to transform NEB 5-alpha chemically competent *E. coli* (NEB C2987H). Twenty-five µL of thawed cells were incubated on ice with 3 µL of the assembly reaction for 30 minutes. After a 30-second incubation at 42 °C and a 5-minute incubation on ice, 1 µL of the transformation reaction was removed for transformant scale determination as described below. The remaining volume of cells was cultured in LB media containing 100 µg/mL carbenicillin (LB_carb_) overnight. The following day, a glycerol stock was prepared by combining 500 µL of culture with 500 µL of filtered 50% glycerol.

### Determination of transformation scale

The transformation reaction prepared above was diluted at 1:20 and 1:100 in LB_carb_ and 20 µL was streaked onto solid media of LB-agar supplemented with 100 µg/mL carbenicillin. For each dilution the colony forming units (CFUs) were determined and used to estimate the number of transformants in the 20 µL used to inoculate the overnight culture, yielding 5,250 cells and a 16-fold oversampling of the 328-member library.

### Initial estimation of library coverage from sampling statistics

The expected fractional coverage of a library (*F*) in each sample is given by

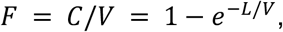

where *C* = unique variants sampled, *V* = total number of unique variants in the library, and *L* = total number of samples taken (e.g., number of transformants or number of sorted wells).^16,17^ The level of “oversampling” (e.g., transformation scale) is represented by the ratio of *L/V* in the exponent. This equation, derived from the Poisson distribution which approximates this sampling process, assumes sampling with replacement and an unbiased distribution of variant abundance in the population.

### Single-cell *E. coli* FACS protocol

Twelve hours before sorting, the *E. coli* pool was used to inoculate a culture containing LB_carb_ and grown with shaking at 37 °C. A 100-µL volume of the saturated cell culture was diluted into 900 µL of a buffer (e.g., 1X phosphate-buffered saline; Thermo 10010023; but similar results were achieved with LB). Cells were washed twice to remove residual media by pelleting via centrifugation at 10,000×*g*, removing the supernatant, and gently resuspending the cells in 1 mL of new buffer. Sorting was performed on either a Sony SH800S or BD FACSAria II instrument, using either a 70 µm or 100 µm nozzle. Similar performance was achieved across all instrument configurations. Prior to sorting the cell sample, a sample of the buffer used for dilution was analyzed to observe the distribution of scattering signal for non-cell events. This was used to gate against the cell sample to reduce the frequency of false positives. Sorted cells were isolated within the wells of transparent-bottom 384-well plates (Thermo 242764) that were pre-filled with 40 µL of LB_carb_ (*Note*: The transparent plate bottom is critical for optical density measurements in the steps to follow). Sorted plates were subjected to centrifugation at 4,000×*g* and then tightly covered with adhesive foil seals (Thermo AB0626) and incubated at 37 °C for 24 hours without shaking.

After 24 hours, single colonies were present at the bottom of wells. (*Note*: at this stage images can be obtained to identify doublets.) Colonies were resuspended via pipetting using a 96-channel Liquidator96 (Rainin 17010335) and 1 µL was used in the subsequent barcoding PCRs (below). Glycerol stock plates were prepared simultaneously by thoroughly mixing 10 µL of the culture with 10 µL of 50% (v/v) glycerol in a second 384-well plate and stored at -80 °C. The OD_600_ of each well was measured by a plate reader (Tecan) and used to determine sorting efficiency.

### IDR library cloning and transformation

The library was ordered as pooled Multiplex Gene Fragments from Twist Biosciences. DNA was assembled into a plasmid destination vector (see **Supporting Information**) by combining 3.38 µL of the library (33.8 ng; 10 ng/µL total library DNA) and 0.625 µL of the destination vector (75 ng; 120 ng/µL total vector DNA) in a 20 µL Golden Gate^30^ reaction using NEBridge BsaI-HFv2 Golden Gate master mix (NEB E1601S). Assembly reactions were then transformed into NEB 5-alpha chemically competent *E. coli* (NEB C2987H). A 50 µL aliquot of cells was incubated on ice with 5 µL of the assembly reaction for 30 minutes. After a 30-second incubation at 42 °C and a 5-minute incubation on ice, 1 µL of the transformation reaction was removed for transformant scale determination as described above. The remaining volume of cells was cultured in LB media containing 100 µg/mL carbenicillin (LB_carb_) overnight. The following day, a glycerol stock was prepared by combining 500 µL of culture with 500 µL of filtered 50% glycerol.

After demultiplexing and picking hits from the library, gene fragments were ordered as eBlocks from Integrated DNA Technologies. Each fragment was assembled into the destination vector by combining 10 ng of fragment and 45 ng of vector DNA in a 10 µL assembly reaction, from each of which 1 µL was transformed into 10 µL of NEB 5-alpha chemically competent *E. coli* (NEB C2987H) following the transformation steps above. Four colonies were picked per construct and sequence verified via the uSort-M sequencing pipeline.

### Optimized uSort-M barcoding protocol

LevSeq-style barcodes were ordered containing a region of homology to the T7 promoter (forward primers) or T7 terminator (reverse primers) at the 3’ end. Upstream of this was a 24-mer barcode, and upstream of this was appended the evSeq universal adapter sequence. This adds an additional set of “buffer” bases and a known, well-behaving adapter sequence if downstream molecular biology is needed. Sequences can be found in **Supporting Information**. Barcode pair plates were prepared by “stamping” 1 µM (final) of the forward barcode (FBC) plate into a solution containing 1 µM (final) of a reverse barcode (RBC) designated for that plate. In other words, all wells in plate 1 contain RBC-A1, wells in plate 2 contain RBC-A2, and so on, for all 96 RBCs. Each well contains its respective FBC (A1 in A1, A2 in A2, etc.). Groups of four 96-well plates were combined into individual 384-well plates according to the strategy described for the initial uSort-M barcoding approach (see **Supporting Information, Initial Barcoding and Demultiplexing Scheme** and **Figure S11**).

A Q5 PCR master mix was prepared (**Supporting Information, Table S10**) and 7 µL of this mix was distributed to each well of a 384-well PCR plate. Reagents were kept on ice until ready to perform the thermal cycle. Using the same pipette tips to reduce cost, the following transfers were made into the PCR plate in the following order: 2 µL of LevSeq-style barcode primers were added to each well, followed by 1 µL of resuspended overnight culture. (Note from the previous section that a glycerol stocking step was usually performed prior to the transfer of the overnight culture, which also resuspends the cells.) PCR was performed with the following thermal cycle, with a variable extension time (‘X’) depending on the length of the amplicon: 2 min, 98 °C; 10 × [10 s, 98 °C; 30 s, 65→55 °C; X s, 72 °C]; 25 × [10 s, 98 °C; 30 s, 66 °C; X s, 72 °C]; 3 min, 72 °C; Hold, 4 °C. The ‘→’ indicates a touchdown annealing step, starting at 65 °C and decreasing 1 °C every cycle, ending at 55 °C. We use 30 seconds per kb for the extension time.

In each row of the barcoding plates, 5 µL from each well was pooled along each row (120 µL total) and added to a 200 µL PCR strip tube containing 30 µL of 100 mM EDTA. From each row-wise pool, 75 µL (1,200 µL total) was pooled into a 1.5 mL tube representing the combined plate. From each plate-wise pool, 200 µL was combined with 130 µL of SPRIselect beads (Beckman Coulter B23318) to perform a left-side selection of >300 bp (although cutoff size can be adjusted to accommodate smaller amplicons). DNA was eluted in 20 µL of DI water and quantified using a Qubit HS kit (Thermo Q33231) at an appropriate dilution (typically 1:100). The resulting library was sequenced via nanopore long-read sequencing with sufficient coverage to yield a depth of >100 reads in each well showing successful outgrowth (see **Supporting Information, Initial Barcoding and Demultiplexing Scheme**).

### Hit picking

#### Clone selection

A map linking each desired variant to its source plate and well position was compiled from the sequencing analysis using the usortm software package, which includes an input file formatted for an Integra ASSIST PLUS pipetting robot (see **Supporting Information** for file format).

#### Liquid handling

Glycerol stock plates were thawed on ice for 30 minutes. Using the ASSIST PLUS, 5 µL of each identified clone was transferred from the source glycerol stock into 100 µL of LB in a destination 96-well plate. Destination plates were incubated overnight at 37 °C with shaking at 250 rpm. Following overnight growth, cultures were mixed 1:1 with 50% filtered glycerol to create 25% glycerol stocks for long-term storage at -80 °C.

For downstream applications, glycerol stocks were used to inoculate expression cultures or subjected to plasmid miniprep for sequence verification.

## Acknowledgements

We thank Forrest L. Zepezauer for assistance with initial cloning experiments, and Michael T. Montgomery and Benjamin R. Doughty for preparing sequencing libraries and performing Illumina sequencing. We thank the Stanford Shared FACS Facility for instrument access and technical support. This work was supported by the National Institute of General Medical Sciences (NIGMS) of the National Institutes of Health (NIH) under grant number R01GM064798; the NIH Office of the Director under grant number 5DP1CA290563; and a National Science Foundation CAREER award under grant number 2142336. Individual support comes from the National Human Genome Research Institute (NHGRI) of the NIH (F31HG013267 to M.B.O.), the National Institute of General Medical Sciences (NIGMS) of the NIH (F32GM156066 to P.J.A.), and the Fannie and John Hertz Foundation, in partnership with Eric and Wendy Schmidt (to L.K.B.). P.M.F. is a Chan Zuckerberg Biohub San Francisco Investigator. The content is solely the responsibility of the authors and does not necessarily represent the official views of the NIH.

## Author Contributions

M.B.O., P.J.A., and P.M.F. conceived the study. M.B.O. and P.J.A. developed the methodology, performed experiments, analyzed data, developed software, and wrote the manuscript with input from all authors. P.J.A. and M.B.O. developed simulations and modeling. L.K.B. designed the IDR library and L.K.B. and M.B.O. implemented the refined workflow method. J.D.S. performed initial feasibility testing for single-cell sorting of *E. coli*. P.M.F. supervised the research, provided funding, and edited the manuscript.

