## Supporting Information for "uSort-M: Scalable isolation of user-defined sequences from diverse pooled libraries"

### Table of Contents

**Supplemental Methods**
1. Minimum cost per variant calculation
2. Initial barcoding and demultiplexing scheme
3. evSeq demultiplexing program call
4. Dorado demultiplexing program calls
5. Abundance estimation for the hAcyP2 input library
6. Integra ASSIST PLUS hit picking input file format

**Supplemental Tables**
Table S1. Costs for hAcyP2 library generation with uSort-M
Table S2. Twist Biosciences Oligonucleotide and Gene Pool pricing tiers
Table S3. Successfully cultured wells after FACS across all eight sorted 384-well plates
Table S4. Nextera i7/i5 barcodes for plate-based indexing
Table S5. Read classification statistics during demultiplexing for MiSeq and Nanopore
Table S6. evSeq PCR mastermix for one 384-well plate
Table S7. Example evSeq refseq files for odd plates
Table S8. Example evSeq refseq files for even plates
Table S9. Fold sampling required to reach 90% library coverage across simulation parameters
Table S10. LevSeq PCR mastermix for one 384-well plate

**Supplemental Figures**
Figure S1. Cost scaling comparison
Figure S2. Construct sequence diagram for hAcyP2 library
Figure S3. Analysis of optical densities of cultured wells after FACS-based isolation
Figure S4. Imaging-based detection of doublet sorts

Figure S5. Analysis of reads assigned to low-OD wells by MiSeq and Nanopore demultiplexing pipelines
Figure S6. Comparison of MiSeq and Nanopore sequencing pipelines as a function of amplicon position
Figure S7. Example well containing multiple variants
Figure S8. Library skew of hAcyP2 input library
Figure S9. Cost and sampling efficiency for exhaustive oversampling vs. targeted resynthesis
Figure S10. Construct sequence diagram for IDR library
Figure S11. Mapping scheme for evSeq and Nextera barcodes for all eight sorted 384-well plates

**DNA Sequences**

1. Wild-type hAcyP2 sequence (codon-optimized for *E. coli*)
2. Oligo pool amplification primers
3. Destination vector sequence
4. Reference sequence for minimap2 alignment

### Supplemental Methods

#### Minimum uSort-M cost per variant calculation

We computed a realistic minimum uSort-M cost per variant (CPV) of $2.58 based on a specific use case: a library of short oligonucleotides (30 bp) encoding single substitutions assembled into a 2,000-bp wild-type gene to generate a library of 2,000 variants. This scenario assumes a 2,000-bp full-length gene where variants are generated by installing short variable regions into a constant backbone sequence via pooled Golden Gate assembly. The site-directed mutagenesis (SDM) comparison assumes per-variant PCR-based mutagenesis of the wild-type backbone with mutagenic primers (~30 nt) and a commercial kit (NEB Q5 SDM, with optional HiFi reassembly), followed by arrayed transformation, consumables, and barcoded pooled sequence validation, with an assumed 10% per-reaction failure rate requiring reclone attempts. Direct synthesis, by comparison, assumes total direct synthesis of each ORF, followed by arrayed cloning and sequence validation by barcoded pooled sequencing. Under these conditions, the minimum reagent cost breakdown per variant is as follows:

| **Step** | **uSort-M** | **SDM** | **Direct** |
|---|---|---|---|
| Synthesis | $0.68 | $15.12 | $105.00 |
| Cloning | $0.09 | $5.36 | $6.12 |
| Sorting | $0.26 | N/A | N/A |
| Barcoding | $0.88 | $0.25 | $0.25 |
| Sequencing | $0.53 | $0.25 | $0.25 |
| Hit picking | $0.14 | N/A | N/A |
| **Total** | $2.58 | $24.06 | $172.60 |
| **Library Total** | $5,409 | $48,127 | $345,199 |

This cost calculation assumes: (1) low library skew (2-fold 90/10 skew; see **Figure S8** for evaluation of Twist Biosciences Oligonucleotide Pool library uniformity), (2) 67% sorting efficiency, (3) low off-target variation (≤10%), (4) 6-fold oversampling during sorting, (5) library resynthesis to recover remaining dropouts, and (6) a target coverage of >90%. The CPV continues to decrease as library size grows beyond 2,000 sequences, but we provide this example as a large but realistic library. For longer gene fragments or libraries with higher complexity, costs will increase proportionally due to higher synthesis costs and greater required oversampling (see **Figure S1** for cost projections across library sizes and fragment lengths).

#### Initial barcoding and demultiplexing scheme

*evSeq and Nextera barcoding PCRs*

Well-based barcoding of the sorted AcyP scanning library followed the evSeq protocol[14] to create the initial, well-specific inner barcoded amplicon via a two-step, one-pot PCR scheme. Briefly, 1 µL of resuspended cell culture from sorted plates was added to 7 µL of evSeq PCR master mix containing site-specific inner primers, Taq polymerase, DMSO, dNTPs, and ThermoPol buffer (**Table S6** and ref. 14). An initial PCR was performed with the following thermal cycle: 5 min, 95 °C; 10 × [20 s, 95 °C; 20 s, 63→53 °C; 30 s, 68 °C]. ('→' indicates a touchdown annealing step, starting at 63 °C and decreasing 1 °C every cycle, ending at 53 °C.)

Following this, well-specific barcodes were added to these reactions according to the following scheme, which is graphically outlined in **Figure S11**: for odd-numbered plates (1, 3, 5, 7), dual-index barcoding plates DI01–DI04 (96 unique barcode pairs each) were used to barcode each well in the plate. The 384-well plates were treated as four unique interleaved 96-well plates, with each one mapping to alternating wells originating from A1, A2, B1, or B2. (For example, A1 wells are the union of columns 1, 3, 5, …, 24 and rows A, C, E, …, P.) DI01 was added to wells associated with A1, DI02 with A2, DI03 with B1, and DI04 with B2. For even-numbered plates (2, 4, 6, 8), DI05–DI08 were used and similarly associated with A1, A2, B1, and B2, respectively. After addition of 2 µL of the DI plate barcodes, the PCR was continued with the following thermal cycle: 25 × [20 s, 95 °C; 20 s, 53 °C; 30 s, 68 °C]; 5 min, 68 °C.

At the completion of the PCR, wells for each unique DI plate were combined into a single mixture containing a final concentration of 20 mM EDTA, pH 8 to inhibit further polymerase activity. (This is performed efficiently by using a multichannel pipette to transfer 5 µL from each well into a strip of 12 PCR tubes pre-filled with 10 µL of 100 mM EDTA, pH 8. Then the mixture from each PCR tube is combined into a single well of a deep-well 96-well plate.) Reactions originating from the same origin 384-well plates were then further combined into microcentrifuge tubes. These eight mixtures were then analyzed and purified by gel electrophoresis using 2% agarose.

Purified DNA was then subjected to a short secondary PCR that appends the plate-specific, outer Nextera barcodes onto each amplicon (**Table S4**). Addition of these barcodes fully differentiates plates that otherwise share the same set of well-specific barcode sets (e.g., odd and even-numbered plates) into fully unique DNA samples that can be pooled and sequenced together.

Pooled DNA from each plate was amplified with KAPA HiFi HotStart polymerase at a scale of 1 µL DNA at a concentration of 1 ng/µL, 12.5 µL of 2X KAPA HiFi HotStart ReadyMix (Roche KK2602), 10 µL of $H_2O$, 0.75 µL of 10 µM forward primer, and 0.75 µL of 10 µM reverse primer (see **DNA Sequences**). Thermal cycling conditions were as follows: 95 °C, 5 min; 10 × [98 °C, 20 s; 60 °C, 15 s; 72 °C, 2 min]; 72 °C, 2 min.

*Illumina MiSeq sequencing and analysis*

Library preparation. Pooled amplicons were purified using Ampure XP beads (Beckman Coulter A63881) at a 1.8x ratio according to the manufacturer's protocol. Purified library concentration was quantified using a Qubit HS Assay Kit (Thermo Fisher Q32854) and fragment size was verified by eGel. The final pooled library concentration was 12.25 ng/µL with an average fragment size of 590 bp.

Sequencing. Libraries were diluted to 4 nM and denatured with 0.2 N NaOH for 5 minutes at room temperature. Denatured libraries were diluted to 9 pM in Illumina Hybridization Buffer and pooled with 30% PhiX Sequencing Control v3 (Illumina FC-110-3001). A 600 µL aliquot of the library was loaded onto a MiSeq Reagent Kit v3 (600-cycle) cartridge (Illumina MS-102-3003) and sequenced with 300 cycles on R1 and 300 cycles on R2.

Basecalling and demultiplexing. Basecalling was performed using bcl2fastq (v2.20) on the Stanford Sherlock HPC cluster. Individual 384-well plates were demultiplexed by their 12-bp Nextera indices (see **Table S4** for barcode sequences), then further demultiplexed to well positions using the evSeq software[14] with default parameters and flags `--keep_parsed_fastqs` and `--return_alignments` (See next sections for exact commands, **Tables S6–7** for example refseq files). Total read depth and composition for each well was determined with the well-specific FASTQ files generated from the previous step.

*Nanopore Long-read sequencing and analysis*

Sample submission. From the Illumina library preparation, 40 µL of pooled amplicons at 6 ng/µL were submitted to Plasmidsaurus for Custom DNA Sequencing with a target yield of 3 Gb (https://www.plasmidsaurus.com/custom).

Basecalling and demultiplexing. Basecalling was performed by Plasmidsaurus using Dorado (v4.3.0), and FASTQ files were downloaded for downstream analysis. Reads were demultiplexed using Oxford Nanopore's Dorado demux tool (v1.1.1; see next sections for parameters). The same 12-bp Nextera and 7-bp evSeq barcode sequences used for Illumina demultiplexing were supplied as a custom barcode arrangement file (see next sections for configuration details).

Variant calling. Demultiplexed reads for each well were aligned to the hAcyP2 reference sequence using minimap2[34] (2.28) with the following command:

```
minimap2 -ax map-ont -A 4 -B 2 -O 10,24 [reference.fasta] [well.fastq] \
-o [well.sam]
```

where bracketed parameters are specific to the data being processed (see **DNA Sequences** for the reference sequence used for `reference.fasta`). Obtained SAM files were converted to BAM files and sorted with the samtools[35] commands: `samtools view -bS [well.sam] | samtools sort -o [well.bam]`. Consensus sequences were generated using samtools consensus (v1.21) with the following command:

```
samtools consensus [well.bam] -aA -o [well.fasta] -c 0.95 -d 100 \
--show-del yes --show-ins yes --mark-ins -q
```

and then combined into a single FASTA file for downstream analysis. Variants calls were made by comparing each consensus sequence to the reference; wells were assigned to library members when ≥90% of reads supported a single variant call. Total read depth and composition for each well were determined from the well-specific FASTQ files.

#### evSeq demultiplexing program call

Using the refseq files shown in **Tables S6–7**, the following code is saved as an executable `.sh` file and run with the provided instructions. The refseq files should be in the same directory and the path to the plate-specific fastq files adjusted.

```
#!/bin/bash

# note: install evSeq environment and run this with `conda run -n evSeq ./run_evSeq.sh`

for i in $(seq 1 8);
do
    refseq="plate${i}refseq.csv"
    data="../MiSeq/fastq_output/plate${i}/"
    evSeq $refseq $data --keep_parsed_fastqs --return_alignments

done
```

#### Dorado demultiplexing program calls

Dorado was used to process the initial multiplexed fastq data, demultiplexing it according to the Nextera barcodes using the following command:

```
dorado demux data.fastq --kit-name nextera_bcs_trim --barcode-
arrangement nextera_bcs_trim.toml --barcode-sequences
nextera_i7rc.fasta --barcode-both-ends --no-trim -o NXT_demux/
```

where the parameter file `nextera_bcs_trim.toml` was:

```
[arrangement]
name = "nextera_bcs_trim"
kit = "CZI"

mask1_front = "AATGATACGGCGACCACCGAGATCTACAC"
mask1_rear = "TCGTCGGCAGCGTC"
mask2_front = "CAAGCAGAAGACGGCATACGAGAT"
mask2_rear = "GTCTCGTGGGCTCGG"

# Barcode sequences
barcode1_pattern = "CZB-NXT-i5-%02i"
barcode2_pattern = "CZB-NXT-i7-%02i"
first_index = 1
last_index = 8

## Scoring options
[scoring]
max_barcode_penalty = 6
min_barcode_penalty_dist = 2
min_separation_only_dist = 5
min_flank_score = 0.8
barcode_end_proximity = 100
front_barcode_window = 175
rear_barcode_window = 175
flank_left_pad = 5
flank_right_pad = 5
midstrand_flank_score = 0.95
```

and the barcode file `nextera_i7rc.fasta` was:

```
>CZB-NXT-i7-01
CCACACAAGAGA
>CZB-NXT-i5-01
TTTAGTCATTGA
>CZB-NXT-i7-02
GTCAGATACCAC
>CZB-NXT-i5-02
AAGATACAAGAG
>CZB-NXT-i7-03
ATAAGCCTTCTG
>CZB-NXT-i5-03
GAGGATACACAT
```

```
>CZB-NXT-i7-04
TGCTGGTGGCTA
>CZB-NXT-i5-04
CCGCCCGCTCTA
>CZB-NXT-i7-05
AGAGGACGAGGA
>CZB-NXT-i5-05
TCCGCTCGGTAA
>CZB-NXT-i7-06
TGGGCATTGTCC
>CZB-NXT-i5-06
CCTCACGCATCG
>CZB-NXT-i7-07
TGGATTGTGATA
>CZB-NXT-i5-07
TCGGACTCCTCG
>CZB-NXT-i7-08
GAAGTTACCACG
>CZB-NXT-i5-08
CCAGCCATCCCG
```

where the i7 indices were the reverse complement of what is provided in Table S4 to match the sequencing orientation. This step demultiplexed the data into nine .bam files: one for reads classified as one of the eight possible barcode pairs (indicated by the final two digits of the sequence names in the fasta file, representing each plate) and one for unclassified reads.

Reads were similarly demultiplexed according to their evSeq barcodes. Each of the eight demultiplexed files from the previous step were subject to four rounds of demultiplexing, one per evSeq dual-index plate (DI01–DI08), according to the following file/DI plate mappings:

```
index_pairs = {
    1: [1, 2, 3, 4],
    2: [5, 6, 7, 8],
    3: [1, 2, 3, 4],
    4: [5, 6, 7, 8],
    5: [1, 2, 3, 4],
    6: [5, 6, 7, 8],
    7: [1, 2, 3, 4],
    8: [5, 6, 7, 8],
}
```

In other words, reads in plate 1 were demultiplexed using evSeq DI plates 1–4, plate 2 using DI 5–8, and so on. Plates demultiplexed using non-target DI plates (e.g., plate 1 using DI05, which should not have barcode pairs present in plate 1) returned only a small number of assigned reads, indicating high accuracy during the demultiplexing process. The command for barcode file (plate) 1 using DI01 was:

```
dorado demux NXT_demux/barcode01.bam --kit-name evseq_bcs --barcode-
arrangement evSeq_bcs.toml --barcode-sequences evSeq_DI01.fasta --
barcode-both-ends --no-trim --emit-fastq -o
NXT_demux/evSeq_plate01_DI01_demux
```

where the parameter file `evseq_bcs.toml` was:

```
[arrangement]
name = "evseq_bcs"
kit = "evSeq"

mask1_front = "TCGTCGGCAGCGTCAGATGTGTATAAGAGACAG"
mask1_rear = "CACCCAAGACCACTCTCCGG"
mask2_front = "GTCTCGTGGGCTCGGAGATGTGTATAAGAGACAG"
mask2_rear = "CGGTGTGCGAAGTAGGTGC"

# Barcode sequences
barcode1_pattern = "evSeq-FBC-%02i"
barcode2_pattern = "evSeq-RBC-%02i"
first_index = 1
last_index = 96

## Scoring options
[scoring]
max_barcode_penalty = 2
min_barcode_penalty_dist = 1
min_separation_only_dist = 3
min_flank_score = 0.7
barcode_end_proximity = 150
front_barcode_window = 150
rear_barcode_window = 150
flank_left_pad = 0
flank_right_pad = 10
midstrand_flank_score = 0.95
```

and the barcode file `evSeq_DI01.fasta` was:

```
>evSeq-FBC-01
GATCATG
>evSeq-RBC-01
GAACTGC
>evSeq-FBC-02
TACATGG
>evSeq-RBC-02
ACCAGGT
…
>evSeq-FBC-96
CCTAATC
>evSeq-RBC-96
ACTCAAC
```

where the number corresponds to the well (A1=01, A2=02, …, H12=96) and the sequences are identical to the provided evSeq barcode sequences for the well in the given plate. A different fasta file was prepared for each of DI01–DI08 matching this pattern and used to further demultiplex each file. The function call was updated and run for each possible plate/DI plate combination. Each unique call generated a new folder (32 total) that contained 97 .fastq files: one for each barcode combination and one for unclassified reads. The unclassified reads were expectedly high, as it included true unclassified and reads that belonged to the three other DI plates.

**Abundance estimation for the hAcyP2 input library**

From the PCR-amplified, gel extracted dsDNA pool used for hAcyP2 library assembly, 5 μL of material at 13.6 ng/μL was submitted to Plasmidsaurus for Premium PCR amplicon sequencing (https://plasmidsaurus.com/amplicon). From the resulting FASTQ file containing 9.3k reads, we filtered for reads in the expected range of 297 ± 10 bp and used minimap2 (v2.28) to map variant reference sequences to reads. Mapped read counts were normalized to the mean of the distribution and library skew was calculated as the ratio of the $90^{th}$ to $10^{th}$ percentile abundance values.

**Input file format for automated hit picking with the Integra ASSIST PLUS robot**

Following sequencing analysis, we generated a hit list mapping each desired variant to its source plate and well position. This hit list was reformatted as an input file compatible with the Integra ASSIST PLUS liquid handling robot to automate cherry-picking of verified clones from glycerol stock plates. The ASSIST PLUS accepts semicolon-delimited CSV files with manufacturer-defined column headers. The required columns are:

| Column | Description |
|---|---|
| SampleID | User-defined identifier for the sample (e.g., variant name) |
| SourcePlateID | Numeric identifier for the source glycerol stock plate (1-indexed) |
| SourceWell | Well position in the source plate (e.g., A1, K23) |
| TargetPlateID | Numeric identifier for the destination plate (0-indexed) |
| TargetWell | Well position in the destination plate |
| TransferVolume | Volume to transfer in μL |

Example input file:

```
SampleID;SourcePlateID;SourceWell;TargetPlateID;TargetWell;TransferVolume
K44A;1;K23;0;G12;5
G45A;1;A11;0;G14;5
T46G;1;G7;0;G15;5
T46A;1;K10;0;G16;5
T48A;1;A21;0;G20;5
V39F;1;E12;0;H1;5...
```

Source plates are numbered sequentially starting from 1, corresponding to the sorted 384-well glycerol stock plates. Destination plates are numbered starting from 0. Well positions use standard alphanumeric notation (rows A–P for 384-well plates, columns 1–24). A Python script for converting sequencing analysis output to ASSIST PLUS-compatible format is available at https://github.com/FordyceLab/usortm. The script accepts a CSV file containing variant names and source well locations, then generates the formatted hit list with user-specified destination plate layouts.

### Supplemental Tables

**Table S1.** Costs for AcyP library generation with uSort-M.

| Item | Manufacturer (Cat #) | Quantity | Units | Cost each | AcyP library total cost |
|---|---|---|---|---|---|
| Oligo pool (101–500 oligos, 251–300 nt) | Twist Bioscience | 1 | pool | $2,000.00 | $2,000.00 |
| DNA assembly reagents (Golden Gate) | NEB (E1601S) | 20 | reactions | $178.00 | $26.70 |
| Chemically competent *E. coli* | NEB (C2987H) | 1 | mL | $215.00 | $5.38 |
| 384-well culture plate | Thermo (242764) | 9 | each | $12.03 | $108.27 |
| Media + antibiotics | | | | | *trivial* |
| FACS instrument rate | | 1.66 | hour | $135 | $225.00 |
| 384-well PCR plate | Bio-Rad (HSP3801) | 8 | each | $7.84 | $62.72 |
| Tips (single PCR) | Rainin (17014404) | 3072 | each | $0.15 | $460.80 |
| dNTPs | NEB (N0447L) | 650 | µL | $0.08 | $48.75 |
| Taq Polymerase (with ThermoPol buffer) | NEB (M0267E) | 160 | µL | $0.35 | $56.48 |
| DMSO (molecular biology grade) | Sigma (D8418) | | | | *trivial* |
| Barcoding primers | IDT | | | | *trivial* |
| Tips (pooling) | Integra 125 µL GripTips (4425) | 384 | each | $0.13 | $49.15 |
| DNA quantification | Qubit HS (Q33231) | | | | *trivial* |
| Sequencing | Plasmidsaurus Custom | 3 | GB | N/A | $600.00 |
| Plates (cherry picking) | Bio-Rad (HSP3801) | 1 | each | $7.84 | $7.84 |
| Tips (cherry picking) | Integra 12.5 µL XYZ Racks (3704) | 384 | each | $0.13 | $49.15 |
| **Total** | | | | | **$3,700.24** |

| ***Cost to barcode each 384-well plate:*** | | | | | |
|---|---|---|---|---|---|
| 384-well culture plate | Thermo (242764) | 1 | each | $12.03 | $12.03 |
| 384-well PCR plate | Bio-Rad (HSP3801) | 1 | each | $7.84 | $7.84 |
| Tips (single PCR) | Rainin (17014404) | 384 | each | $0.15 | $57.60 |
| dNTPs | NEB (N0447L) | 80 | µL | $0.08 | $6.00 |
| Taq Polymerase (with ThermoPol buffer) | NEB (M0267E) | 20 | µL | $0.35 | $7.06 |
| Tips (pooling) | Integra 125 µL GripTips (4425) | 48 | each | $0.15 | $7.20 |
| **Total** | | | | | **$97.73** |

**Table S2**. Twist Biosciences Oligonucleotide Pool (top) and Gene Pool (bottom) pricing tiers as of May 2026.

| **Tier (Pool size)** | 20–120nt | 121–150nt | 151–200nt | 201–250nt | 251–300nt | 301–350nt |
|---|---|---|---|---|---|---|
| Tier 1 (2–100 Oligos) | $400.00 | $466.00 | $520.00 | $689.00 | $1,030.00 | $1,288.00 |
| Tier 2 (101–500 Oligos) | $800.00 | $933.00 | $1,040.00 | $1,379.00 | $2,060.00 | $2,575.00 |
| Tier 3 (501–1000 Oligos) | $1,200.00 | $1,400.00 | $1,560.00 | $2,068.00 | $3,090.00 | $3,863.00 |
| Tier 4 (1001–2000 Oligos) | $1,600.00 | $1,867.00 | $2,080.00 | $2,757.00 | $4,121.00 | $5,152.00 |
| Tier 5 (2001–6000 Oligos) | $2,400.00 | $2,800.00 | $3,120.00 | $4,136.00 | $6,181.00 | $7,727.00 |
| Tier 6 (6001–12000 Oligos) | $3,120.00 | $3,744.00 | $4,056.00 | $5,148.00 | $7,694.00 | $9,618.00 |
| Tier 7 (12001–18000 Oligos) | $4,056.00 | $4,867.00 | $5,273.00 | $6,694.00 | $10,004.00 | $12,505.00 |
| Tier 8 (18001–24000 Oligos) | $5,273.00 | $6,327.00 | $6,855.00 | $8,702.00 | $13,006.00 | $16,258.00 |

| **Tier (Pool size)** | ≤650bp | 651–1,400bp |
|---|---|---|
| Tier 1 (2–1,000 Genes) | $17,335.00 | $19,000.00 |
| Tier 2 (1,001–2,000 Genes) | $24,660.00 | $32,058.00 |
| Tier 3 (2,001–6,000 Genes) | $39,299.00 | $51,089.00 |
| Tier 4 (6,001–12,000 Genes) | $64,680.00 | $84,084.00 |
| Tier 5 (12,001–18,000 Genes) | $87,120.00 | $113,256.00 |
| Tier 6 (18,001–24,000 Genes) | $108,240.00 | $140,712.00 |

**Table S3.** Successfully cultured wells (determined by optical density) after FACS across all eight sorted 384-well plates.

| Plate | Wells | Percent (%) |
|---|---:|---:|
| 1 | 264 | 68.8 |
| 2 | 248 | 64.6 |
| 3 | 267 | 69.5 |
| 4 | 263 | 68.5 |
| 5 | 251 | 65.4 |
| 6 | 265 | 69.0 |
| 7 | 241 | 62.8 |
| 8 | 258 | 67.2 |
| Total | 2,057 | 67.0 |

**Table S4.** Nextera i7/i5 barcodes for plate-based indexing.

| Plate | Index 1 (i7) | Index 2 (i5) |
|---|---|---|
| 1 | TCTCTTGTGTGG | TTTAGTCATTGA |
| 2 | GTGGTATCTGAC | AAGATACAAGAG |
| 3 | CAGAAGGCTTAT | GAGGATACACAT |
| 4 | TAGCCACCAGCA | CCGCCCGCTCTA |
| 5 | TCCTCGTCCTCT | TCCGCTCGGTAA |
| 6 | GGACAATGCCCA | CCTCACGCATCG |
| 7 | TATCACAATCCA | TCGGACTCCTCG |
| 8 | CGTGGTAACTTC | CCAGCCATCCCG |

**Table S5.** Read classification statistics during the demultiplexing processes for MiSeq and Nanopore sequencing data with the AcyP library. [a]Includes expected unclassified reads from PhiX DNA addition to compensate for the low-diversity amplicon library.

| Step | MiSeq | Nanopore |
|---|---|---|
| Total Reads | 16,387,889 | 4,307,220 |
| Unclassified Outer | 10,843,812[a] | 2,029,862 |
| Total Inner Reads | 5,544,077 | 2,277,358 |
| Total Inner Assigned to Wells | 3,446,033 | 1,467,668 |
| Fraction Inner Assigned to Wells | 0.622 | 0.644 |
| Wells with >100 reads | 2,017 | 2,005 |

**Table S6.** evSeq PCR master mix for one 384-well plate.

| Component | Amount (µL) |
|---|---|
| Thermopol Buffer | 576 |
| 10 mM dNTPs | 115.2 |
| Taq Polymerase | 28.8 |
| Water | 3070 |
| DMSO | 230.4 |
| Inner Primer Mix (10 µM each) | 11.52 |

**Table S7.** Example evSeq refseq files for odd plates. Only the plate number in the PlateName column is changed for different refseq files for odd plates.

| PlateName | IndexPlate | FPrimer | RPrimer | VariableRegion | FrameDistance | BpIndStart | AaIndStart |
|---|---|---|---|---|---|---|---|
| Plate1_DI01 | DI01 | CACCCAAGA<br>CCACTCTCC<br>GGTGGCCCA<br>TGAAGGTCA<br>TCGTC | CGGTGTGCG<br>AAGTAGGTG<br>CTGAACAGC<br>TCCTCGCCC | TGGGTAAACCGGGTGAGGGACCTGCATCTGGAGGTGGA<br>AGCGGAGGAAGTGGAGGTGGAAGCACCGCGCAGAGCCT<br>GAAAAGCGTGGATTATGAAGTGTTTGGCCGCGTGCAGG<br>GCGTGTGCTTTCGCATGTATACCGAAGATGAAGCGCGC<br>AAAATTGGCGTGGTGGGCTGGGTGAAAAACACCAGCAA<br>AGGCACCGTGACCGGCCAGGTGCAGGGCCCGGAAGATA<br>AAGTGAACAGCATGAAAAGCTGGCTGAGCAAAGTGGGC<br>AGCCCGAGCAGCCGCATTGATCGCACCAACTTTAGCAA<br>CGAAAAAACCATTAGCAAACTGGAATATAGCAACTTTA<br>GCATTCGCTATGGTGGCGGTTCAGGGTCGGGCGGTTCA<br>GGGTCTGTTAGCAA | 2 | 19 | 8 |
| Plate1_DI02 | DI02 | CACCCAAGA<br>CCACTCTCC<br>GGTGGCCCA<br>TGAAGGTCA<br>TCGTC | CGGTGTGCG<br>AAGTAGGTG<br>CTGAACAGC<br>TCCTCGCCC | TGGGTAAACCGGGTGAGGGACCTGCATCTGGAGGTGGA<br>AGCGGAGGAAGTGGAGGTGGAAGCACCGCGCAGAGCCT<br>GAAAAGCGTGGATTATGAAGTGTTTGGCCGCGTGCAGG<br>GCGTGTGCTTTCGCATGTATACCGAAGATGAAGCGCGC<br>AAAATTGGCGTGGTGGGCTGGGTGAAAAACACCAGCAA<br>AGGCACCGTGACCGGCCAGGTGCAGGGCCCGGAAGATA<br>AAGTGAACAGCATGAAAAGCTGGCTGAGCAAAGTGGGC<br>AGCCCGAGCAGCCGCATTGATCGCACCAACTTTAGCAA<br>CGAAAAAACCATTAGCAAACTGGAATATAGCAACTTTA<br>GCATTCGCTATGGTGGCGGTTCAGGGTCGGGCGGTTCA<br>GGGTCTGTTAGCAA | 2 | 19 | 8 |
| Plate1_DI03 | DI03 | CACCCAAGA<br>CCACTCTCC<br>GGTGGCCCA<br>TGAAGGTCA<br>TCGTC | CGGTGTGCG<br>AAGTAGGTG<br>CTGAACAGC<br>TCCTCGCCC | TGGGTAAACCGGGTGAGGGACCTGCATCTGGAGGTGGA<br>AGCGGAGGAAGTGGAGGTGGAAGCACCGCGCAGAGCCT<br>GAAAAGCGTGGATTATGAAGTGTTTGGCCGCGTGCAGG<br>GCGTGTGCTTTCGCATGTATACCGAAGATGAAGCGCGC<br>AAAATTGGCGTGGTGGGCTGGGTGAAAAACACCAGCAA<br>AGGCACCGTGACCGGCCAGGTGCAGGGCCCGGAAGATA<br>AAGTGAACAGCATGAAAAGCTGGCTGAGCAAAGTGGGC<br>AGCCCGAGCAGCCGCATTGATCGCACCAACTTTAGCAA<br>CGAAAAAACCATTAGCAAACTGGAATATAGCAACTTTA<br>GCATTCGCTATGGTGGCGGTTCAGGGTCGGGCGGTTCA<br>GGGTCTGTTAGCAA | 2 | 19 | 8 |
| Plate1_DI04 | DI04 | CACCCAAGA<br>CCACTCTCC<br>GGTGGCCCA<br>TGAAGGTCA<br>TCGTC | CGGTGTGCG<br>AAGTAGGTG<br>CTGAACAGC<br>TCCTCGCCC | TGGGTAAACCGGGTGAGGGACCTGCATCTGGAGGTGGA<br>AGCGGAGGAAGTGGAGGTGGAAGCACCGCGCAGAGCCT<br>GAAAAGCGTGGATTATGAAGTGTTTGGCCGCGTGCAGG<br>GCGTGTGCTTTCGCATGTATACCGAAGATGAAGCGCGC<br>AAAATTGGCGTGGTGGGCTGGGTGAAAAACACCAGCAA<br>AGGCACCGTGACCGGCCAGGTGCAGGGCCCGGAAGATA<br>AAGTGAACAGCATGAAAAGCTGGCTGAGCAAAGTGGGC<br>AGCCCGAGCAGCCGCATTGATCGCACCAACTTTAGCAA<br>CGAAAAAACCATTAGCAAACTGGAATATAGCAACTTTA<br>GCATTCGCTATGGTGGCGGTTCAGGGTCGGGCGGTTCA<br>GGGTCTGTTAGCAA | 2 | 19 | 8 |

**Table S8.** Example evSeq refseq files for even plates. Only the plate number in the PlateName column is changed for different refseq files for even plates.

| PlateName | IndexPlate | FPrimer | RPrimer | VariableRegion | FrameDistance | BpIndStart | AaIndStart |
|---|---|---|---|---|---|---|---|
| Plate2_DI05 | DI05 | CACCCAAGACCACTCTCCGGTGGCCCATGAAGGTCATCGTC | CGGTGTGCGAAGTAGGTGCTGAACAGCTCCTCGCCC | TGGGTAAACCGGGTGAGGGACCTGCATCTGGAGGTGGAAGCGGAGGAAGTGGAGGTGGAAGCACCGCGCAGAGCCTGAAAAGCGTGGATTATGAAGTGTTTGGCCGCGTGCAGGGCGTGTGCTTTCGCATGTATACCGAAGATGAAGCGCGCAAAATTGGCGTGGTGGGCTGGGTGAAAAACACCAGCAAAGGCACCGTGACCGGCCAGGTGCAGGGCCCGGAAGATAAAGTGAACAGCATGAAAAGCTGGCTGAGCAAAGTGGGCAGCCCGAGCAGCCGCATTGATCGCACCAACTTTAGCAACGAAAAAACCATTAGCAAACTGGAATATAGCAACTTTAGCATTCGCTATGGTGGCGGTTCAGGGTCGGGCGGTTCAGGGTCTGTTAGCAA | 2 | 19 | 8 |
| Plate2_DI06 | DI06 | CACCCAAGACCACTCTCCGGTGGCCCATGAAGGTCATCGTC | CGGTGTGCGAAGTAGGTGCTGAACAGCTCCTCGCCC | TGGGTAAACCGGGTGAGGGACCTGCATCTGGAGGTGGAAGCGGAGGAAGTGGAGGTGGAAGCACCGCGCAGAGCCTGAAAAGCGTGGATTATGAAGTGTTTGGCCGCGTGCAGGGCGTGTGCTTTCGCATGTATACCGAAGATGAAGCGCGCAAAATTGGCGTGGTGGGCTGGGTGAAAAACACCAGCAAAGGCACCGTGACCGGCCAGGTGCAGGGCCCGGAAGATAAAGTGAACAGCATGAAAAGCTGGCTGAGCAAAGTGGGCAGCCCGAGCAGCCGCATTGATCGCACCAACTTTAGCAACGAAAAAACCATTAGCAAACTGGAATATAGCAACTTTAGCATTCGCTATGGTGGCGGTTCAGGGTCGGGCGGTTCAGGGTCTGTTAGCAA | 2 | 19 | 8 |
| Plate2_DI07 | DI07 | CACCCAAGACCACTCTCCGGTGGCCCATGAAGGTCATCGTC | CGGTGTGCGAAGTAGGTGCTGAACAGCTCCTCGCCC | TGGGTAAACCGGGTGAGGGACCTGCATCTGGAGGTGGAAGCGGAGGAAGTGGAGGTGGAAGCACCGCGCAGAGCCTGAAAAGCGTGGATTATGAAGTGTTTGGCCGCGTGCAGGGCGTGTGCTTTCGCATGTATACCGAAGATGAAGCGCGCAAAATTGGCGTGGTGGGCTGGGTGAAAAACACCAGCAAAGGCACCGTGACCGGCCAGGTGCAGGGCCCGGAAGATAAAGTGAACAGCATGAAAAGCTGGCTGAGCAAAGTGGGCAGCCCGAGCAGCCGCATTGATCGCACCAACTTTAGCAACGAAAAAACCATTAGCAAACTGGAATATAGCAACTTTAGCATTCGCTATGGTGGCGGTTCAGGGTCGGGCGGTTCAGGGTCTGTTAGCAA | 2 | 19 | 8 |
| Plate2_DI08 | DI08 | CACCCAAGACCACTCTCCGGTGGCCCATGAAGGTCATCGTC | CGGTGTGCGAAGTAGGTGCTGAACAGCTCCTCGCCC | TGGGTAAACCGGGTGAGGGACCTGCATCTGGAGGTGGAAGCGGAGGAAGTGGAGGTGGAAGCACCGCGCAGAGCCTGAAAAGCGTGGATTATGAAGTGTTTGGCCGCGTGCAGGGCGTGTGCTTTCGCATGTATACCGAAGATGAAGCGCGCAAAATTGGCGTGGTGGGCTGGGTGAAAAACACCAGCAAAGGCACCGTGACCGGCCAGGTGCAGGGCCCGGAAGATAAAGTGAACAGCATGAAAAGCTGGCTGAGCAAAGTGGGCAGCCCGAGCAGCCGCATTGATCGCACCAACTTTAGCAACGAAAAAACCATTAGCAAACTGGAATATAGCAACTTTAGCATTCGCTATGGTGGCGGTTCAGGGTCGGGCGGTTCAGGGTCTGTTAGCAA | 2 | 19 | 8 |

**Table S9**. Fold sampling required to reach 90% library coverage across simulation parameters. Monte Carlo simulations performed with 1000-member library, holding all other parameters at default optima. N/A = conditions where 90% coverage was not reached within 250-fold sampling.

| Parameter | Value | Fold Sampling at 90% |
|---|---|---|
| Library Skew | 1 | 2.4 |
| Library Skew | 2 | 2.592 |
| Library Skew | 4 | 3.456 |
| Library Skew | 10 | 5.76 |
| Library Skew | 50 | 27.648 |
| Library Skew | 100 | 115.2 |
| Transformation Scale | 1 | N/A |
| Transformation Scale | 2.5 | 6.528 |
| Transformation Scale | 5 | 3.072 |
| Transformation Scale | 10 | 2.688 |
| Transformation Scale | 25 | 2.496 |
| Transformation Scale | 50 | 2.4 |
| Fraction Incorrect | 0 | 2.4 |
| Fraction Incorrect | 0.1 | 2.688 |
| Fraction Incorrect | 0.2 | 2.976 |
| Fraction Incorrect | 0.3 | 3.456 |
| Fraction Incorrect | 0.4 | 4.224 |
| Fraction Incorrect | 0.5 | 4.992 |

**Table S10.** **L**evSeq-style PCR master mix for one 384-well plate (5% excess).

| **Component** | **Amount (µL)** |
|---|---:|
| Q5 5X Buffer | 806.4 |
| 10 mM dNTPs | 80.64 |
| Q5 Polymerase | 40.32 |
| Water | 1733.76 |
| DMSO | 161.28 |

### Supplemental Figures

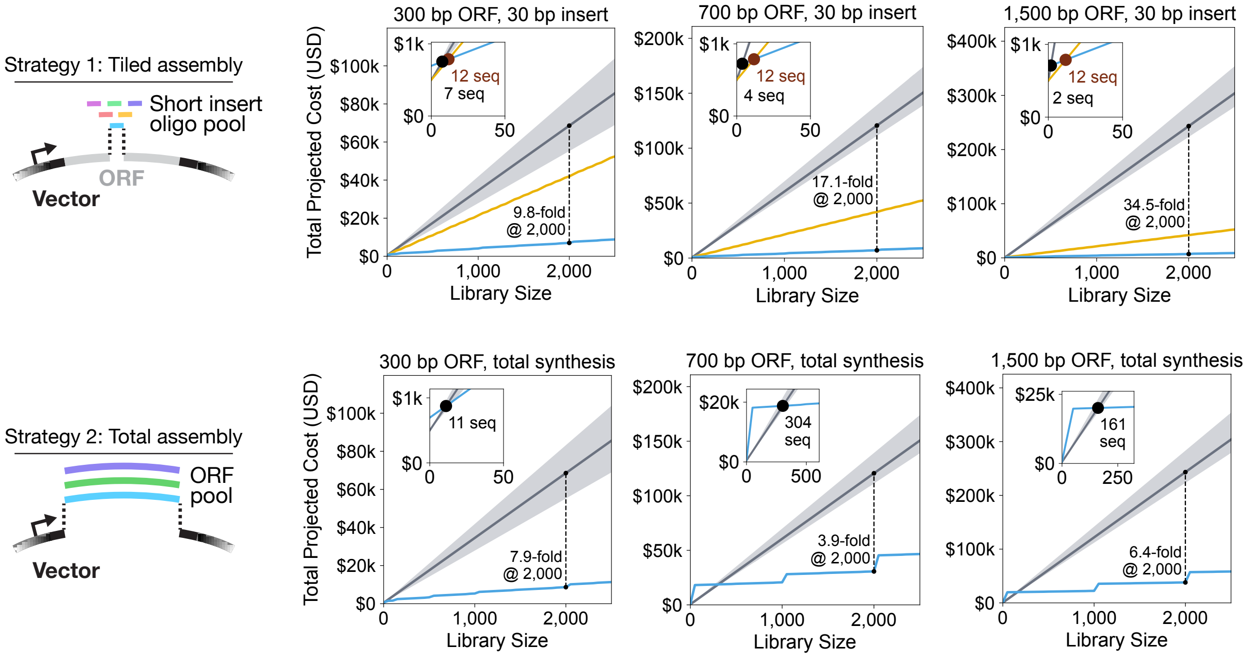


**Figure S1. Cost scaling comparison**. Total projected cost as a function of library size across three ORF lengths (300, 700, 1,500 bp) and two uSort-M assembly strategies, comparing uSort-M (blue), site-directed mutagenesis (SDM, yellow), and direct gene synthesis (grey line; shaded grey region spans the minimum and maximum across representative commercial platforms). Annotations within each inset mark the library sizes at which uSort-M costs cross those of direct synthesis (black point) and SDM (auburn point). Dashed vertical lines indicate a 2,000-variant library size; the adjacent label reports the cost-fold savings of uSort-M relative to direct synthesis at that size. **Top row**: library assembly starting from a pool of 30 bp oligos into a destination vector containing the remaining ORF sequence. **Bottom row**: assembly of full-length ORFs into a destination vector. The 300 bp total-synthesis simulation assumes Twist Bioscience Oligo Pool pricing; the 700 bp and 1,500 bp total-synthesis simulations assume Twist Bioscience Gene Pool pricing (see **Table S2** for pricing information). uSort-M curves use a fold-sampling pinned to 90% expected coverage at 4× library skew.

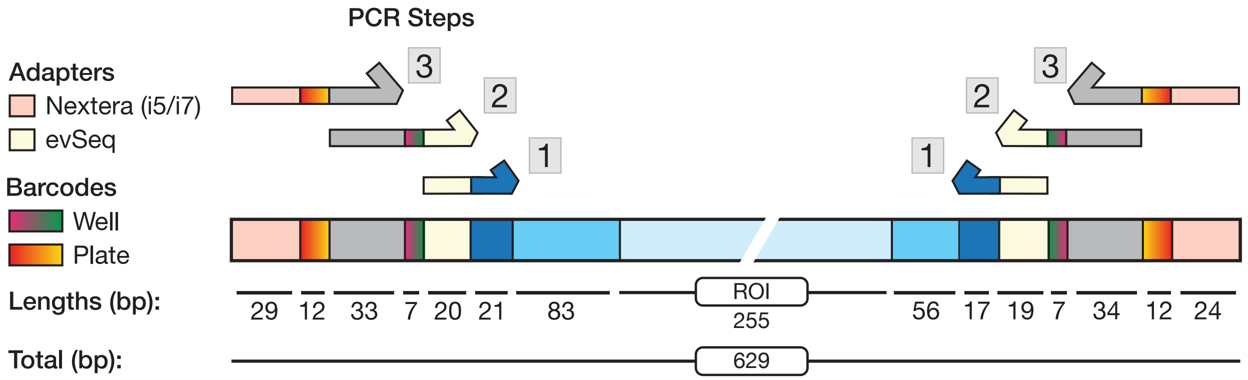


**Figure S2. Construct sequence diagram**. Schematic showing locations and lengths of coding sequences, barcode regions, adapters, and primers used during PCR indexing for the hAcyP2 library. From left to right: pink, Nextera adapter; red/yellow gradient, i5 index (plate-specific); gray, Nextera Read1; purple/green gradient, evSeq FBC index (well-specific); beige, evSeq adapter; dark blue, site-specific evSeq forward priming sequence; medium blue, N-terminal constant region; light blue, region of interest (ROI) containing target variability (i.e., codons 9–93 of hAcyP2, see **DNA Sequences**); medium blue, C-terminal constant region; dark blue, site-specific evSeq reverse priming sequence; beige, evSeq adapter; purple/green gradient, evSeq RBC index (well-specific); gray, Nextera Read2; red/yellow gradient, i7 index (plate-specific); pink, Nextera adapter. All blue regions match DNA in the template plasmid and can include true on- or off-target variability except for the dark blue region, which originates from primer in the final amplicon, not template. All other regions are appended DNA with no expected variability. Primers for PCR step 1 can be found in **DNA Sequences**. Primers and barcodes for PCR step 2 are from the evSeq method. Indices for the Nextera primers used for PCR step 3 can be found in **Table S4**.

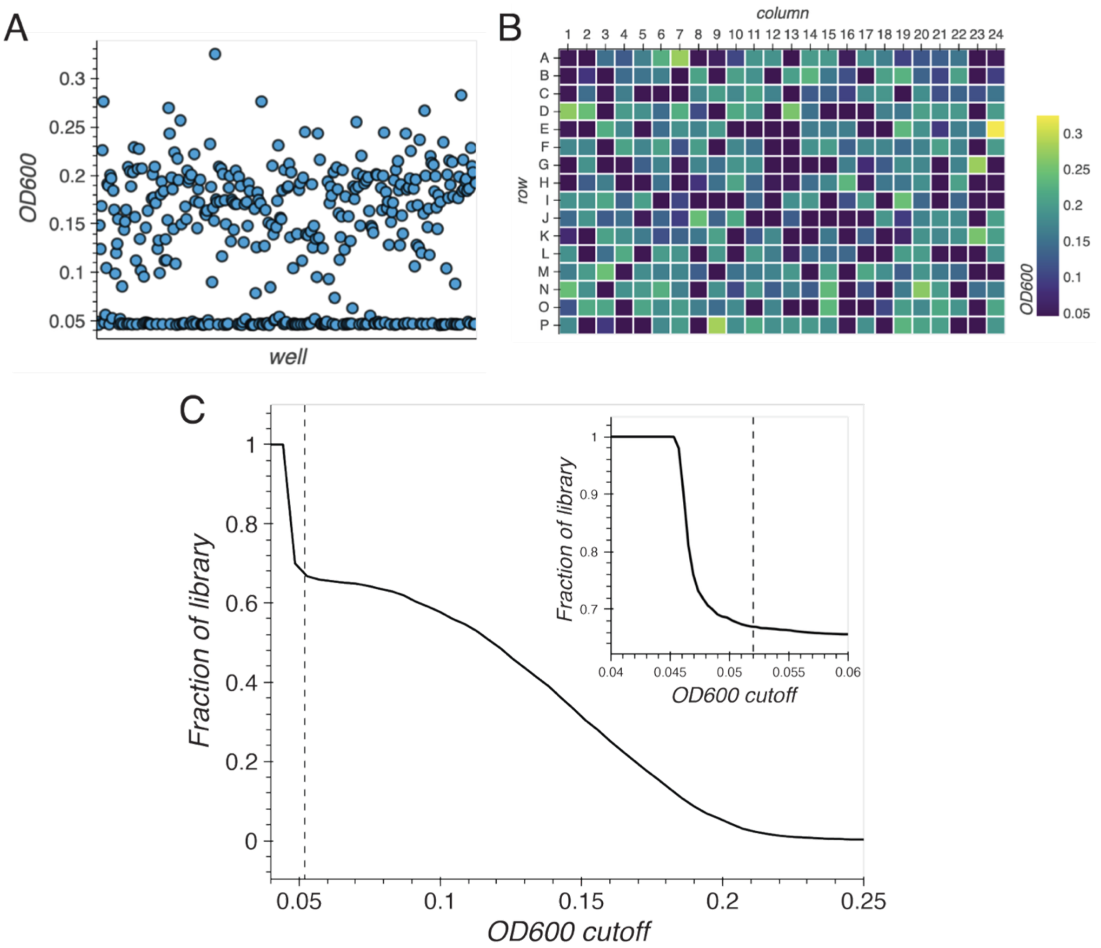


**Figure S3. Analysis of optical densities of cultured wells after FACS-based isolation of single cells. A**. Scatter plot (sorted by well) of optical density at 600 nm ($OD_{600}$) readings for a 384-well plate of cultured cells after overnight anaerobic growth and resuspension. **B**. A heatmap colored by $OD_{600}$ for each well. **C**. Relationship between fraction of wells considered to have growth and the $OD_{600}$ cutoff value used to threshold wells for growth or not. A threshold of 0.052 was selected for this experiment. *Inset*: Same plot with adjusted axes.

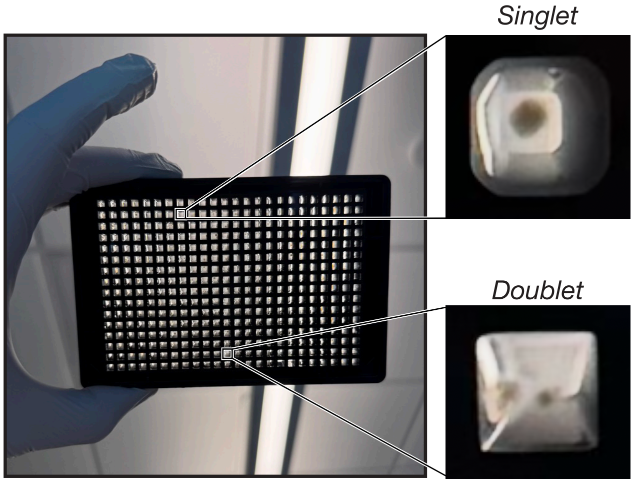


**Figure S4. Image-based detection of doublet sorts**. *Left:* Photo of a sorted plate after 12-hour outgrowth step. *Right*: Zoomed-in images of selected wells. Colonies are visible by eye such that doublet colonies can be readily detected by imaging.

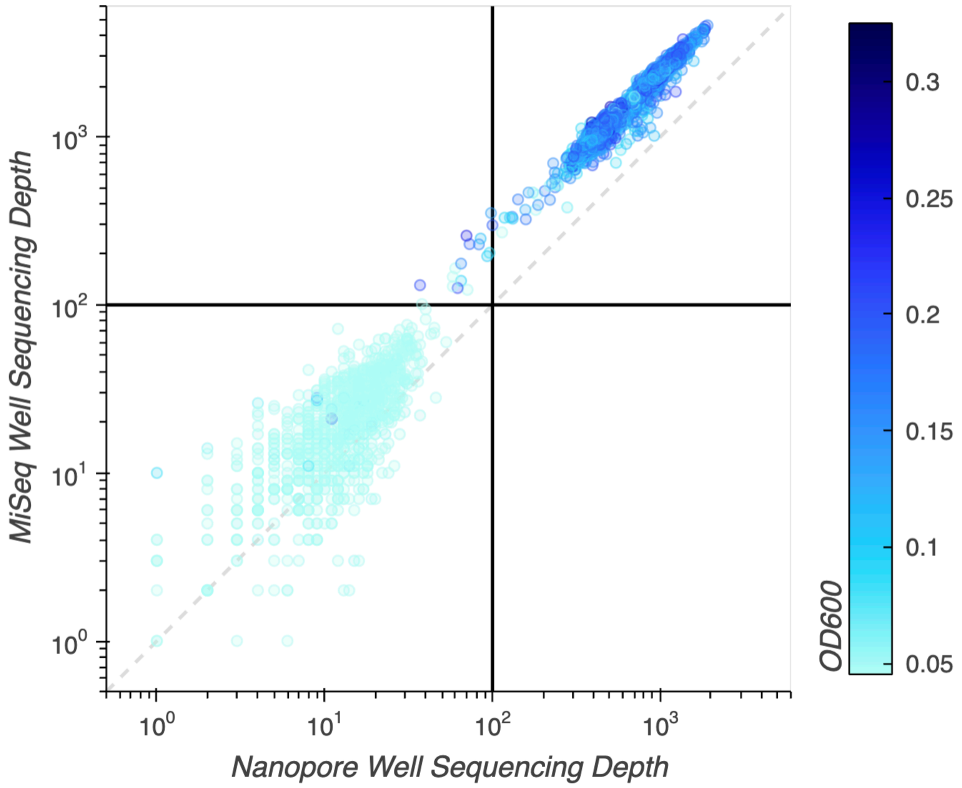


**Figure S5. Analysis of reads assigned to low-OD wells by MiSeq and Nanopore demultiplexing pipelines.** Scatter plot of MiSeq well sequencing depth vs. Nanopore well sequencing depth on a log-log scale. Points are colored by $OD_{600}$; horizontal and vertical lines indicate the 100-read marks for each. Dashed gray line indicates the 1:1 line. No differences are observed for wells with low $OD_{600}$ between either method, which are largely contained within the bottom left quadrant (low read depth).

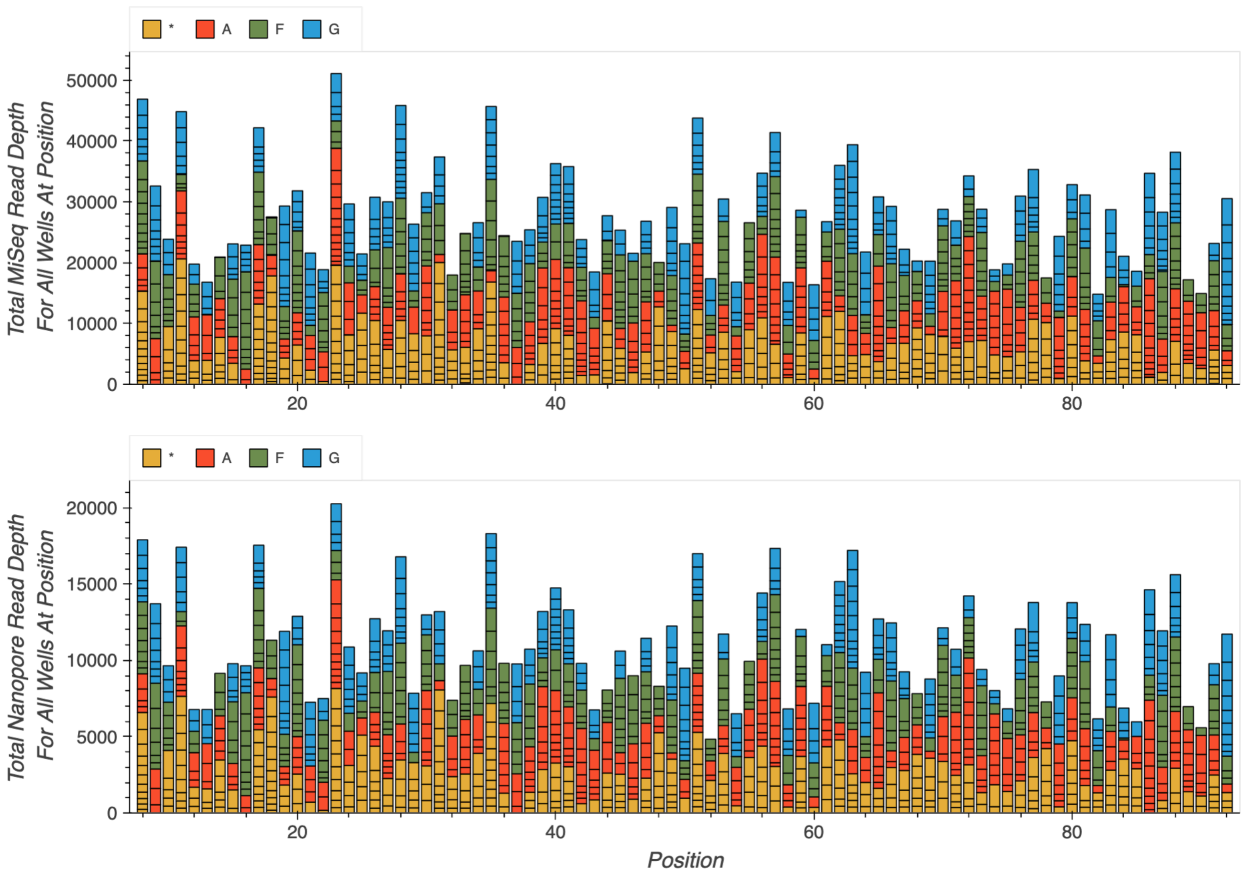


**Figure S6. Comparison of MiSeq (*upper*) and Nanopore (*lower*) sequencing pipelines as a function of amplicon position of the called variant.** A stacked bar is shown for each position in the hAcyP2 amino acid sequence, with the total height representing the summed read depth across all wells that were mapped to that position. Within each stack, the sub-bars convey information about a single well: the height of each sub-bar represents its individual read depth (the number of reads assigned to that specific well) and the color of the sub-bar represents the determined amino acid identity. Sub-bars within each stack are further organized by amino acid identity. The composition of substitutions at each position reflects both library design (WT residues are excluded) and stochastic sampling. Both MiSeq and Nanopore approaches match qualitatively, with no apparent positional bias in variant recovery.

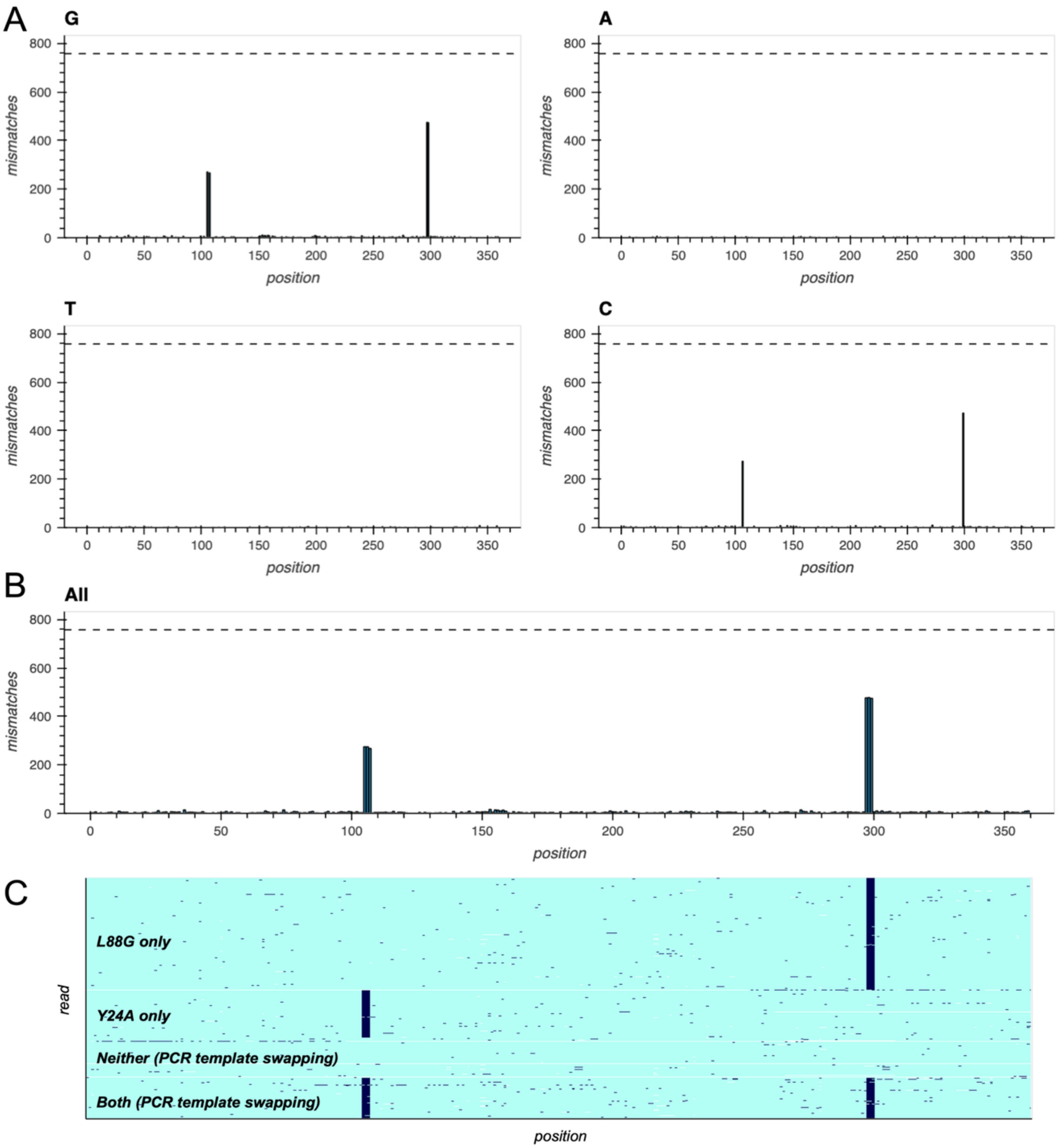


**Figure S7. Example well containing multiple variants**. Representative Nanopore-based read alignments for well A3 from plate 1 (384-well plate nomenclature; 96-well plate nomenclature is plate 1 DI01-A02) showing data from all 758 detected reads. **A**. Base-specific mismatch plots, where the x-axis shows base position and the y-axis shows the sum of identified mismatches that map to G, A, T, or C at each position. **B**. Aggregated mismatch plot, with the same data as shown in (A) but for all bases. **C**. Read-specific mismatch data, which provides information on mismatches that occur within the same read. Reference-matching positions are shown in light blue and mismatches are shown in dark blue; unmapped positions (i.e., from indels) are shown in white. Reads are ordered from top to bottom based on whether they have only the major mutation (corresponding to L88G), only the minor mutation (Y24A), neither, or both. These data most likely indicate that L88G and Y24A are the primary variants in a mixed population, with reads containing neither mutation or both mutations stemming from template swapping between the amplicons generated from these variants during the well-specific barcoding PCR.

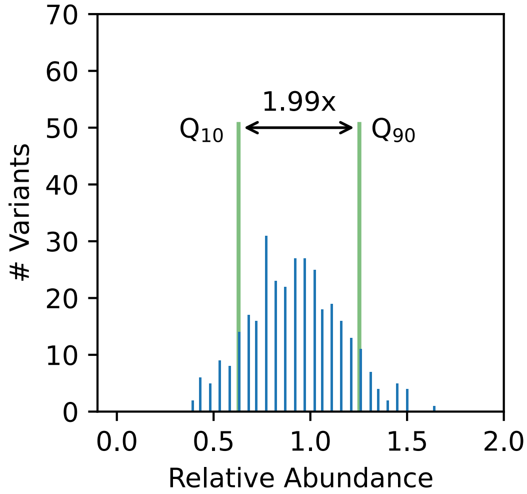


**Figure S8. Skew of hAcyP2 input library**. Histogram of relative abundance for variants within the hAcyP2 input library after initial library amplification. The x-axis shows mean-normalized abundance and y-axis shows the number of unique variants within each abundance bin. Green vertical lines indicate the $10^{th}$ and $90^{th}$ percentiles of the distribution, from which we compute a 10-90 skew of 1.99-fold.

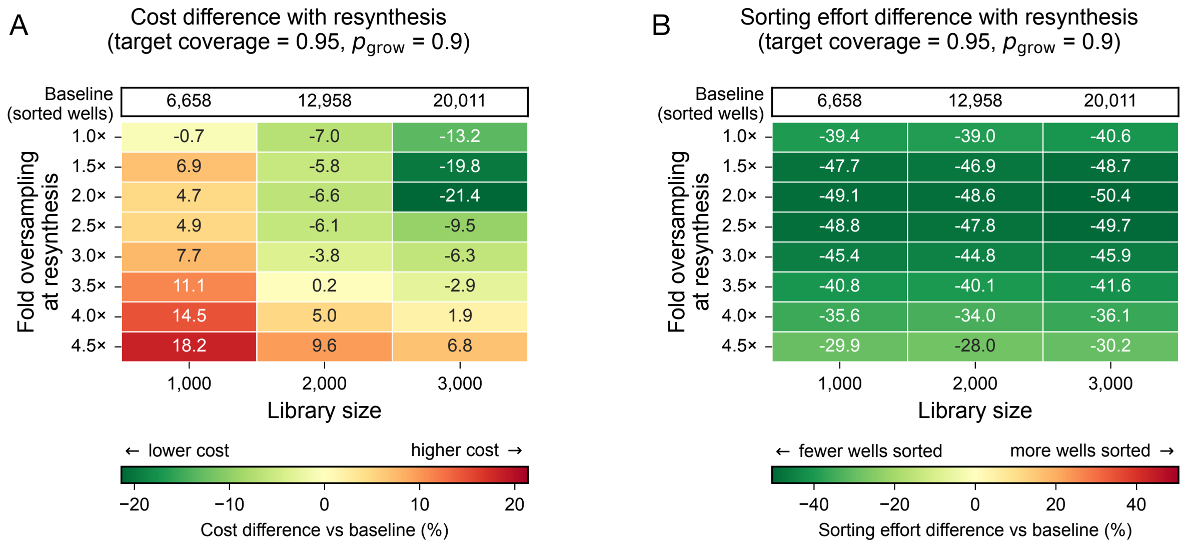


**Figure S9. Cost and sampling efficiency for exhaustive oversampling vs. targeted resynthesis**. **A**. Percentage cost savings as a function of library size and fold-sampling at which resynthesis is triggered. **B**. Reduction in total wells sampled to reach a target coverage of 95% when using iterative resampling compared to single-pass sampling. For both panels, simulations assume 95% target coverage and 90% sorting efficiency; values represent the median of 10 replicate simulations.

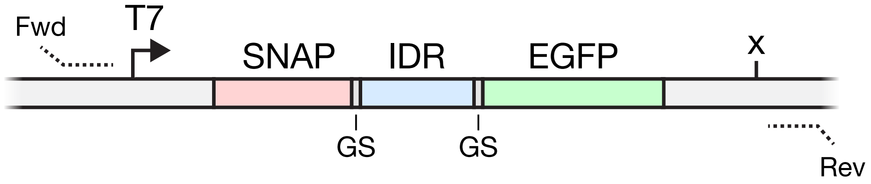


**Figure S10. Construct sequence diagram for IDR library.** Each IDR CDS was assembled into an expression vector containing an N-terminal SNAP tag and C-terminal EGFP tag, connected by glycine-serine linkers (GS). Primer barcodes flanking the T7 promoter and terminator (represented by an “x”) allowed for sequencing of the entire barcoded cassette within a single amplicon.

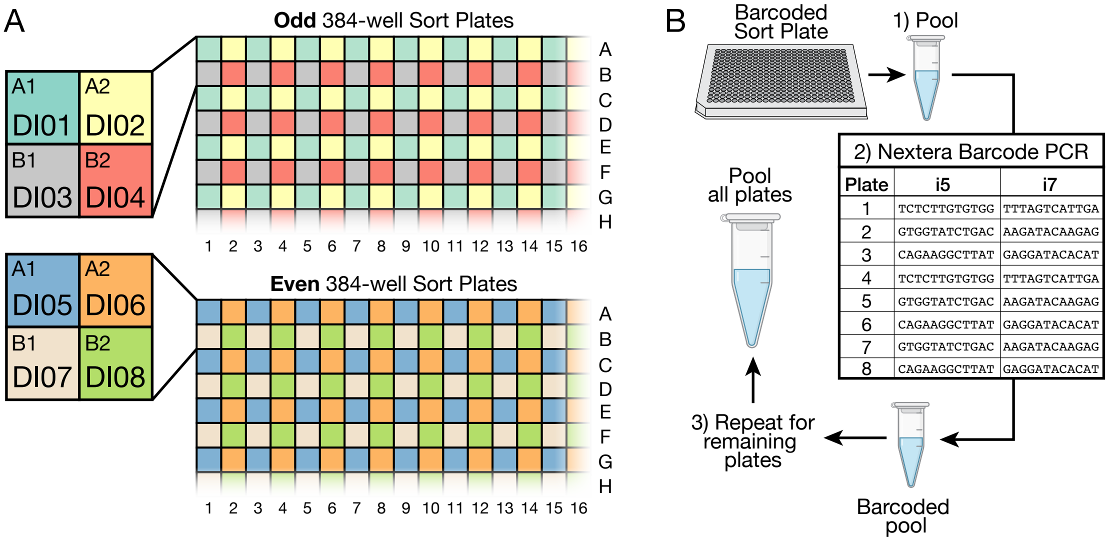


| Plate | i5 | i7 |
|---|---|---|
| 1 | TCTCTTGTGTGG | TTTAGTCATTGA |
| 2 | GTGGTATCTGAC | AAGATACAAGAG |
| 3 | CAGAAGGCTTAT | GAGGATACACAT |
| 4 | TCTCTTGTGTGG | TTTAGTCATTGA |
| 5 | GTGGTATCTGAC | AAGATACAAGAG |
| 6 | CAGAAGGCTTAT | GAGGATACACAT |
| 7 | GTGGTATCTGAC | AAGATACAAGAG |
| 8 | CAGAAGGCTTAT | GAGGATACACAT |

**Figure S11. Mapping scheme for evSeq and Nextera barcodes for all eight sorted 384-well plates. A**. evSeq barcodes used for each plate. Each 96-well plate from evSeq barcode plates DI01–DI08 were mapped to a quadrant of the 384-well sort plates, with plates DI01–DI04 corresponding to odd plates and DI05–DI08 to even plates. **B**. Pooling scheme and Nextera barcoding across 384-well plates. Sort plates were pooled following evSeq barcoding and barcoded with Nextera i5/i7 barcode pairs.

### DNA Sequences

The wild-type hAcyP2 sequence codon-optimized for *E. coli* is:

```
AGCACCGCGCAGAGCCTGAAAAGCGTGGATTATGAAGTGTTTGGCCGCGTGCAGGGCGTGTGCTTT
CGCATGTATACCGAAGATGAAGCGCGCAAAATTGGCGTGGTGGGCTGGGTGAAAAACACCAGCAAA
GGCACCGTGACCGGCCAGGTGCAGGGCCCGGAAGATAAAGTGAACAGCATGAAAAGCTGGCTGAGC
AAAGTGGGCAGCCCGAGCAGCCGCATTGATCGCACCAACTTTAGCAACGAAAAAACCATTAGCAAA
CTGGAATATAGCAACTTTAGCATTCGCTAT
```

To create the hAcyP2 scanning library, all codons from position 9 to 93 were converted to Gly (`GGC`), Ala (`GCG`), Phe (`TTT`), or a stop codon (`TAG`) unless already present in the wild-type sequence, with flanking sequences for library amplification and Golden Gate cloning. All ordered sequences were mutated from the following gene, with the uppercase region corresponding to the variable codons 7–98 of hAcyP2:

```
ggcgcggtctccaaaagcGTGGATTATGAAGTGTTTGGCCGCGTGCAGGGCGTGTGCTTTCGCATG
TATACCGAAGATGAAGCGCGCAAAATTGGCGTGGTGGGCTGGGTGAAAAACACCAGCAAAGGCACC
GTGACCGGCCAGGTGCAGGGCCCGGAAGATAAAGTGAACAGCATGAAAAGCTGGCTGAGCAAAGTG
GGCAGCCCGAGCAGCCGCATTGATCGCACCAACTTTAGCAACGAAAAAACCATTAGCAAACTGGAA
TATAGCAACtttagcattcgctatcctctggcggcg
```

The primers used to amplify the oligo pool were:

Forward: `GGCGCGGTCTCC`

Reverse: `CGCCGCCAGAGG`

The destination vector sequence used for Golden Gate cloning was:

```
GAAGATCCTTTGATCTTTTCTACGGGGTCTGACGCTCAGTGGAACGAAAACTCACAGATCCGGGAT
TTTGGTCATGAGATTATCAAAAAGGATCTTCACCTAGATCCTTTTAAATTAAAAATGAAGTTTTAA
ATCAATCTAAAGTATATATGAGTAAACTTGGTCTGACAGTTACCAATGCTTAATCAGTGAGGCACC
TATCTCAGCGATCTGTCTATTTCGTTCATCCATAGTTGCCTGACTCCCCGTCGTGTAGATAACTAC
GATACGGGAGGGCTTACCATCTGGCCCCAGTGCTGCAATGATACCGCGACTTCCACGCTCACCGGC
TCCAGATTTATCAGCAATAAACCAGCCAGCCGGAAGGGCCGAGCGCAGAAGTGGTCCTGCAACTTT
ATCCGCCTCCATCCAGTCTATTAATTGTTGCCGGGAAGCTAGAGTAAGTAGTTCGCCAGTTAATAG
TTTGCGCAACGTTGTTGCCATCGCTACAGGCATCGTGGTGTCACGCTCGTCGTTTGGTATGGCTTC
ATTCAGCTCCGGTTCCCAACGATCAAGGCGAGTTACATGATCCCCCATGTTGTGCAAAAAAGCGGT
TAGCTCCTTCGGTCCTCCGATCGTTGTCAGAAGTAAGTTGGCCGCAGTGTTATCACTCATGGTTAT
GGCAGCACTGCATAATTCTCTTACTGTCATGCCATCCGTAAGATGCTTTTCTGTGACTGGTGAGTA
CTCAACCAAGTCATTCTGAGAATAGTGTATGCGGCGACCGAGTTGCTCTTGCCCGGCGTCAATACG
GGATAATACCGCGCCACATAGCAGAACTTTAAAAGTGCTCATCATTGGAAAACGTTCTTCGGGGCG
AAAACTCTCAAGGATCTTACCGCTGTTGAGATCCAGTTCGATGTAACCCACTCGTGCACCCAACTG
ATCTTCAGCATCTTTTACTTTCACCAGCGTTTCTGGGTGAGCAAAAACAGGAAGGCAAAATGCCGC
AAAAAAGGGAATAAGGGCGACACGGAAATGTTGAATACTCATACTCTTCCTTTTTCAATATTATTG
AAGCATTTATCAGGGTTATTGTCTCATGAGCGGATACATATTTGAATGTATTTAGAAAAATAAACA
AATAGGGGTTCCGCGCACATTTCCCCGAAAAGTGCTAGTGGTGCTAGCCCCGCGAAATTAATACGA
CTCACTATAGGGTCTAGAAATAATTTTGTTTAACTTTAAGAAGGAGATATACATATGGACAAAGAT
TGCGAAATGAAACGTACCACCCTGGATAGCCCGCTGGGCAAACTGGAACTGAGCGGCTGCGAACAG
GGCCTGCATGAAATTAAACTGCTGGGTAAAGGCACCAGCGCGGCCGATGCGGTTGAAGTTCCGGCC
CCGGCCGCCGTGCTGGGTGGTCCGGAACCGCTGATGCAGGCGACCGCGTGGCTGAACGCGTATTTT
```

CATCAGCCGGAAGCGATTGAAGAATTTCCGGTTCCGGCGCTGCATCATCCGGTGTTTCAGCAGGAG
AGCTTTACCCGTCAGGTGCTGTGGAAACTGCTGAAAGTGGTTAAATTTGGCGAAGTGATTAGCTAT
CAGCAGCTGGCGGCCCTGGCGGGTAATCCGGCGGCCACCGCCGCCGTTAAAACCGCGCTGAGCGGT
AACCCGGTGCCGATTCTGATTCCGTGCCATCGTGTGGTTAGCTCTAGCGGTGCGGTTGGCGGTTAT
GAAGGTGGTCTGGCGGTGAAAGAGTGGCTGCTGGCCCATGAAGGTCATCGTCTGGGTAAACCGGGT
GAGGGACCTGCATCTGGAGGTGGAAGCGGAGGAAGTGGAGGTGGAAGCACCGCGCAGAGCCTGAAA
AAGAGACCATATATATATGGTCTCACTATGGTGGCGGTTCAGGGTCGGGCGGTTCAGGGTCTGTTA
GCAAGGGCGAGGAGCTGTTCACCGGGGTGGTGCCCATCCTGGTCGAGCTGGACGGCGACGTAAACG
GCCACAAGTTCAGCGTGTCCGGCGAGGGCGAGGGCGATGCCACCTACGGCAAGCTGACCCTGAAGT
TCATCTGCACCACCGGCAAGCTGCCCGTGCCCTGGCCCACCCTCGTGACCACCCTGACCTACGGCG
TGCAGTGCTTCAGCCGCTACCCCGACCACATGAAGCAGCACGACTTCTTCAAGTCCGCCATGCCCG
AAGGCTACGTCCAGGAGCGCACCATCTTCTTCAAGGACGACGGCAACTACAAGACCCGCGCCGAGG
TGAAGTTCGAGGGCGACACCCTGGTGAACCGCATCGAGCTGAAGGGCATCGACTTCAAGGAGGACG
GCAACATCCTGGGGCACAAGCTGGAGTACAACTACAACAGCCACAACGTCTATATCATGGCCGACA
AGCAGAAGAACGGCATCAAGGTGAACTTCAAGATCCGCCACAACATCGAGGACGGCAGCGTGCAGC
TCGCCGACCACTACCAGCAGAACACCCCCATCGGCGACGGCCCCGTGCTGCTGCCCGACAACCACT
ACCTGAGCACCCAGTCCGCCCTGAGCAAAGACCCCAACGAGAAGCGCGATCACATGGTCCTGCTGG
AGTTCGTGACCGCTGCCGGGATCACTCTCGGCATGGACGAGCTGTACAAATAATAATGAGGATCCC
GGGAATTCTCGAGTAAGGTTAACCTGCAGGAGGCCTTTAATTAAGGTGGTGCGGCCGCGCTAGCGG
TCCCGGGGGATCGATCCGGCTGCTAACAAAGCCCGAAAGGAAGCTGAGTTGGCTGCTGCCACCGCT
GAGCAATAACTAGCATAACCCCTTGGGGCCTCTAAACGGGTCTTGAGGGGTTTTTTGCTGAAAGGA
GGAACTATATCCGGAAGCTTGGCACTGGCCGACCGGGGTCGAGCACTGACTCGCTGCGCTCGGTCG
TTCGGCTGCGGCGAGCGGTATCAGCTCACTCAAAGGCGGTAATACGGTTATCCACAGAATCAGGGG
ATAACGCAGGAAAGAACATGTGAGCAAAAGGCCAGCAAAAGGCCAGGAACCGTAAAAAGGCCGCGT
TGCTGGCGTTTTTCCATAGGCTCCGCCCCCCTGACGAGCATCACAAAAATCGACGCTCAAGTCAGA
GGTGGCGAAACCCGACAGGACTATAAAGATACCAGGCGTTTCCCCCTGGAAGCTCCCTCGTGCGCT
CTCCTGTTCCGACCCTGCCGCTTACCGGATACCTGTCCGCCTTTCTCCCTTCGGGAAGCGTGGCGC
TTTCTCATAGCTCACGCTGTAGGTATCTCAGTTCGGTGTAGGTCGTTCGCTCCAAGCTGGGCTGTG
TGCACGAACCCCCCGTTCAGCCCGACCGCTGCGCCTTATCCGGTAACTATCGTCTTGAGTCCAACC
CGCTAAGACACGACTTATCGCCACTGGCAGCAGCCACTGGTAACAGGATTAGCAGAGCGAGGTATG
TAGGCGGTGCTACAGAGTTCTTGAAGTGGTGGCCTAACTACGGCTACACTAGAAGAACAGTATTTG
GTATCTGCGCTCTGCTGAAGCCAGTTACCTTCGGAAAAAGAGTTGGTAGCTCTTGATCCGGCAAAC
AAACCACCGCTGGTAGCGGTGGTTTTTTTGTTTGCAAGCAGCAGATTACGCGCAGAAAAAAAGGAT
CTCAA

The site-specific primers used to perform evSeq were:

Forward: CACCCAAGACCACTCTCCGGTGGCCCATGAAGGTCATCGTC

Reverse: CGGTGTGCGAAGTAGGTGCTGAACAGCTCCTCGCCC

The reference sequence used for minimap2 alignment was:

AATGATACGGCGACCACCGAGATCTACACNNNNNNNNNNNNNTCGTCGGCAGCGTCAGATGTGTATA
AGAGACAGNNNNNNNCACCCAAGACCACTCTCCGGTGGCCCATGAAGGTCATCGTCTGGGTAAACC
GGGTGAGGGACCTGCATCTGGAGGTGGAAGCGGAGGAAGTGGAGGTGGAAGCACCGCGCAGAGCCT
GAAAAGCGTGGATTATGAAGTGTTTGGCCGCGTGCAGGGCGTGTGCTTTCGCATGTATACCGAAGA
TGAAGCGCGCAAAATTGGCGTGGTGGGCTGGGTGAAAAACACCAGCAAAGGCACCGTGACCGGCCA
GGTGCAGGGCCCGGAAGATAAAGTGAACAGCATGAAAAGCTGGCTGAGCAAAGTGGGCAGCCCGAG
CAGCCGCATTGATCGCACCAACTTTAGCAACGAAAAAACCATTAGCAAACTGGAATATAGCAACTT
TAGCATTCGCTATGGTGGCGGTTCAGGGTCGGGCGGTTCAGGGTCTGTTAGCAAGGGCGAGGAGCT

```
GTTCAGCACCTACTTCGCACACCGNNNNNNNCTGTCTCTTATACACATCTCCGAGCCCACGAGACN
NNNNNNNNNNNATCTCGTATGCCGTCTTCTGCTTG
```

with the `N`s representing the 12-mer and 7-mer Nextera and evSeq barcodes, and regions matching the schematic in **Figure S2**.
